# Status and future perspectives for pikeperch (*Sander lucioperca*) stocks in Europe

**DOI:** 10.1101/2022.12.20.521162

**Authors:** Eglė Jakubavičiūtė, Timo Arula, Justas Dainys, J. Tyrell Deweber, Harry Gorfine, Laura S. Härkönen, Pekka Hyvärinen, Kristiina Hommik, Jan Kubecka, Linas Ložys, Noora Mustamäki, Rahmat Naddafi, Mikko Olin, Žilvinas Pūtys, Elor Sepp, Allan T. Souza, Andrius Šiaulys, Väino Vaino, Asta Audzijonyte

## Abstract

Pikeperch (*Sander lucioperca*) is a European fresh and brackish water piscivorous fish, important as both a key predator and a valuable commercial and recreational fisheries species. There are concerns that some stocks are depleted due to overfishing and environmental changes. We review data collection and population assessments currently used for nine pikeperch stocks across six European countries and apply a unified assessment framework to evaluate population status and trends. For this we first standardised commercial, scientific, and recreational catch-per-unit-effort (CPUE) and catch time series and then applied Bayesian surplus production models. Our results showed that three stocks (including two in the Baltic Sea) were strongly depleted, with estimated biomasses considerably lower than the biomass at maximum sustainable yield (B_msy_). Other stocks were either close or higher than their estimated B_msy_. Looking at the trends, we find that four stocks (Lake Oulujärvi, Kvädöfjärden, Lake Peipsi and Lipno) showed increasing biomass trends and two (Curonian Lagoon, Galtfjärden) had a strong decline in biomass. In most cases the stocks with clear signs of recovery were also those for which strong management strategies have been implemented. We find that, despite pikeperch being one of the most valuable inland fisheries, formalised stock assessments and regular surveys remain rare. Importantly, although most stocks are strongly targeted by recreational fishing, estimates of recreational catch are highly uncertain. We conclude that data limited stock assessment methods are useful for assessing fish population status and highlight an urgent need to improve pikeperch scientific monitoring and assessment of recreational catches.

## Introduction

Inland and coastal fisheries are important to local communities (Natale et al. 2013; Lynch et al. 2016; Funge-Smith 2018) and are estimated to contribute 12.7% of global capture production (FAO 2022a). At the same time, 70% of European inland basins are under medium to high pressure on a multi-criteria threat scale (FAO 2022a). Inland ecosystem and freshwater fish species are among the most threatened by human impacts, pollution, habitat degradation and climate change (Reid et al. 2019). Yet, most inland fisheries are also data poor or data limited and therefore rarely assessed with common quantitative stock assessments, such as statistical catch-at-age or virtual population analysis. As a result, the status of many stocks remains almost entirely unknown and unassessed, which increases the possibility of stock depletion or inadequate management measures (Hilborn et al. 2020).

Pikeperch (*Sander lucioperca)* is a native species from the family Percidae or ray-finned fishes, common in regions of Europe and Asia and also been introduced into Africa (Welcomme 1988) and North America (Robins et al. 1991). Like other large predatory fishes, it is highly valued by both commercial fisheries and recreational anglers (Raat 1991; Arlinghaus and Mehner 2004; Dainys et al. 2022a). Total pikeperch catch declined from 50 thousand tons in 1950 (FAO 2019) to only 14.6 thousand in 2009, as a result of overfishing (Colchen et al. 2020; FAO 2022b). Since then, the catches have increased to nearly 26 thousand tons in 2020 (FAO 2022b), but this increase needs to be considered in the context of recent range expansions and development of fishing techniques. Moreover, officially reported catches might not reflect true harvest trends, because in most populations recreational fishing may be the main source of fishing mortality, yet very few countries collect regular data on recreational fishing catches and their impacts. For example, a recent detailed assessment in Lithuania showed that recreational catches can greatly exceed commercial catches and impede population recovery after a commercial fisheries closure (Dainys et al. 2022a, b); recreational catches also exceed commercial ones in the Archipelago Sea region of the Baltic Sea (Heikinheimo et al. 2015).

In addition to the importance of commercial and recreational fishing, pikeperch is also one of the main predatory species in European inland and coastal ecosystems, playing a key ecological and regulatory role (Keskinen and Marjomäki 2004; Ranåker et al. 2014). Changes in overall pikeperch abundance and size structure are likely to influence entire ecosystems (Brabrand and Faafeng 1993; Vainikka et al. 2017a). In some intensively harvested populations, fishing has induced decreases in size and age at maturation, suggesting ongoing fishery-induced evolution in its population structure and functional role (Kokkonen et al. 2015), although the true extent of potential life-history changes and evolutionary impact remains largely unassessed. Finally, pikeperch is a warm temperature tolerant species that thrives in warm, turbid, well-oxygenated and productive ecosystems (Sonesten 1991; Veneranta et al. 2011; Froese and Pauly 2022). Due to eutrophication and climate change, environmental conditions for this species may become more suitable across much of central and northern Europe (Müller-Karulis et al. 2013).

Despite its importance commercially, recreationally, and ecologically, current data collection and assessment of European pikeperch populations remains limited. Although some stocks in Finland, Estonia and Sweden have been assessed with age-based virtual population analyses (Eero 2004; Vainikka and Hyvärinen 2012; Heikinheimo et al. 2014), assessments of most stocks are based on mean size trends (Mustamäki et al. 2014), reported commercial or recreational catches (Andrašūnas et al. 2022) or are not undertaken at all (see Table S1). Previous analyses suggest that trends in European pikeperch population status vary. For example, in Finland’s inland waters, growing commercial and recreational catches indicate improved population status (LUKE 2022a, b), likely attributed to increased minimum landing size limits and stocking, as well as global warming and eutrophication. A similar increase in population abundance appears to be occurring in coastal Finnish waters (LUKE 2022a, b). In contrast, fishery independent monitoring of Swedish coastal pikeperch populations in the northeast region of the Baltic Sea suggests large declines in abundance since 1995, associated with heavy commercial fishing and increased great cormorant predation (Mustamäki et al. 2014). Substantial decrease in pikeperch abundance since the end of 1990s is also seen for Pärnu Bay, Estonia, which is due to increased fishing pressure on the stock (Eero 2004; Eschbaum et al. 2021). In contrast, in Lake Peipsi (Estonia), there has been a strong increase in the abundance of pikeperch since the end of the 1980s, mostly due to the limitations imposed on the small-mesh Danish seine fishery (Saat et al. 2010). In Lithuania natural populations occur only in the Nemunas river, Curonian Lagoon and the Baltic Sea (Virbickas 2000), but the species has been stocked into multiple lakes and water reservoirs. A recent catch-only based analysis suggested overexploitation in the Curonian Lagoon (Andrašūnas et al. 2022). Scientific surveys also show that many other populations in the Baltic region have been declining (Bergström et al. 2016). In Kaunas reservoir in Lithuania commercial fishing has been banned since 2013, but a recent study demonstrated a limited recovery rate, which could be explained by large recreational catches that greatly exceed previous commercial catch (Dainys et al. 2022a). In the Czech Republic, pikeperch is an important recreational fisheries species with catches ranging between 50 to 95 tons (Czech Anglers Union 2022). A population established in Lipno Reservoir is one of the most recreationally important and abundant ones. The population, however, showed a strong decline in the early 2000’s, which prompted introductions of new management practices that appears to have improved its status (Vehanen at al. 2020). Pikeperch occurs throughout Upper Lake Constance, but we only use data from Austria where the most suitable habitat is located and where most targeted fishing occurs. The pikeperch fishery in Lake Constance is relatively unimportant compared to other species, and very little is known about stock status or trends.

Lack of uniform and regular assessments of pikeperch in inland and coastal ecosystems is likely to be impeding appropriate management measures because there is good evidence that stock status and sustainability are strongly and positively correlated with management intensity and that fisheries without regular time-varying assessments and management plans are more likely to be depleted or depleting (Melnychuk et al. 2021). Regular assessments are particularly important in light of rapidly changing environmental conditions and increasing popularity and efficiency of recreational fishing, for which catches often remain unknown. In this study we explored pikeperch status in Europe and collected data from recreational, commercial and scientific surveys and known catch records. With these data we statistically standardised the catch per unit effort (CPUE) time series and applied a unified analysis using Bayesian surplus production modelling (Winker et al. 2018). We also reviewed current data collection programs and provide recommendations for improving data collections and monitoring.

## Materials and methods

### Study areas and data

We analysed nine pikeperch populations from six northern and central European countries (Finland, Sweden, Estonia, Lithuania, Czech Republic and Austria), representing a cross section of inland and coastal water bodies with different management strategies (Fig. 1, Table 1). Data used in this study were contributed by the relevant scientists from each country and included commercial effort and catches, recreational catches (where available) as well as scientific surveys (Table S2). We also considered data for the Pärnu Bay population in the Baltic Sea (Estonia), but due to the short CPUE time series (only 10 years) did not include it here.

**Fig. 1.**
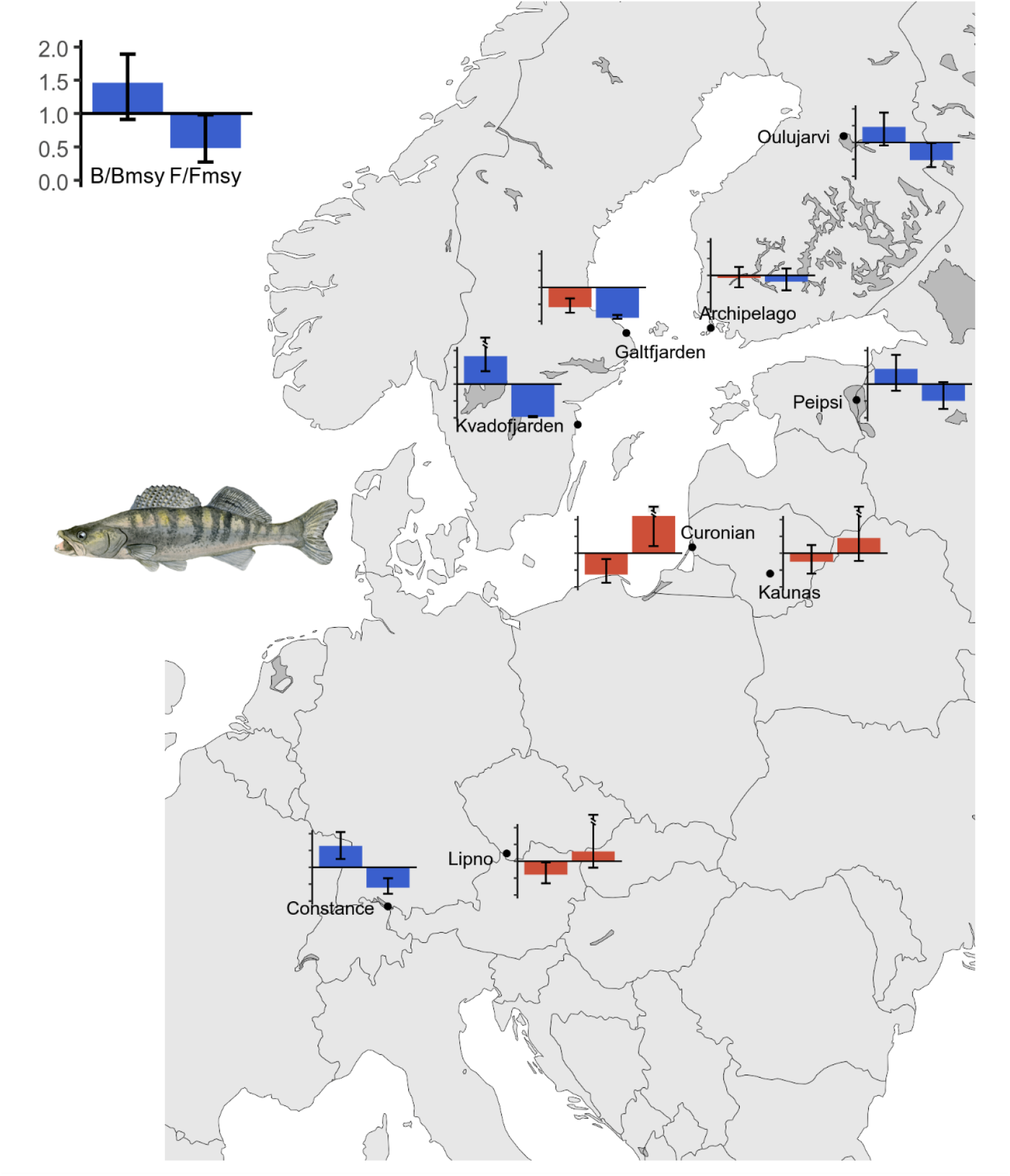
Status of pikeperch stocks (last year in the time series, 2019 to 2021) in the European populations studied. The first bar illustrates the ratio of current biomass (B) versus biomass required for maximum sustainable yield (B_msy_), where values >1 indicate a good status (shown in blue) and values <1 show bad status (red). The second bar shows current fishing mortality (F) versus mortality leading to maximum sustainable yield (F_msy_) and values <1 indicate good status. Error bars show uncertainty intervals from the surplus production models (see Fig. 2).

**Table 1.** Study areas and pikeperch populations included in this study, and their key characteristics and management measures used.

| Study area | Environmental conditions | Commercial harvest management | Recreational harvest management | General trends, threats and other factors | Stocking |
| --- | --- | --- | --- | --- | --- |
| 1. Archipelago Sea, Baltic Sea, Finland | Brackish (7‰), mean depth ~ 23 m, inner bays, area of ca 8300 km<sup>2</sup>, population native | Minimum size limit: TL ≥40 cm | Minimum size limit: TL ≥42 cm; gear limits: 8 gillnets, or in total of 240 m | Commercial catches and effort declining; increasing eutrophication and water temperatures | Yes, few |
| 2. Lake Oulujärvi, Finland | Freshwater, water level regulated; area 778-994 km<sup>2</sup>, shallow (~ 8 m); mesotrophic, native stock collapsed, new stock established in 1985 | Minimum size limit: TL ≥45 cm; gear limits: 2 gillnets with knot size of 20-49 mm, number of gillnets with smaller or larger knot sizes not limited | Minimum size limit: TL ≥45 cm; gear limits: 8 gillnets, or in total of 240 m | Commercial catches relatively stable, increasing water temperature (enhances natural reproduction) | Yes, heavily (~ 300,000 yoy per year) |
| 3. Galtfjärden Bay, Baltic Sea, Sweden | Brackish, shallow (~ 6 m), ca 75 km<sup>2</sup>, population native | Minimum size limit: > 45 cm, also maximum size limit: <60 cm when fishing with hand gear and fyke nets | Minimum size limit: TL >45 cm, also maximum size limit: <60 cm when fishing with hand gear and fyke nets | Predation from seals and cormorants, commercial and recreational fisheries | No |
| 4. Kvädöfjärden, Baltic Sea, Sweden | Brackish (6-8 ‰), shallow (~ 7 m), ca 12 km<sup>2</sup>, population native | Minimum size limit: > 45 cm, also maximum size limit: <60 cm when fishing with hand gear and fyke nets | Minimum size limit: TL >45 cm, maximum size limit: TL <60 cm when fishing with hand gear and fyke nets | Predation from seals and cormorants, commercial and recreational fisheries | No |
| 5. Lake Peipsi, Estonia | Freshwater, shallow (~ 7 m), 3555 km<sup>2</sup>, productive, population native | TAC, gear number, minimum mesh sizes and minimum landing size (≥ 46cm), for Danish seine fishery temporary reduction in MLS), seasonal closures | Minimum size limit: TL >46 cm; daily bag limit of 5 pikeperch since 2021 (recreational catches are negligible) | Important commercial fishery shared between Estonia and Russia | No |
| 6. Curonian Lagoon, Lithuania | Brackish-fresh (0-3‰), shallow (~4 m), 1619 km<sup>2</sup>, highly productive, population native | TAC and TAE until 2013; TAE only from 2014; minimum mesh sizes for gillnets (40 cm), landing size (TL 46 cm), seasonal and spatial closures | Minimum size limit: TL ≥45 cm; daily bag limit of 2 pikeperch (total bag ≤7 kg), seasonal closures (March 1- May 31) | Commercial catches stable, rapidly declining scientific CPUE; most commercial catches taken at the border with Russia; recreational catch likely large, but unknown | No |
| 7. Kaunas Water Reservoir, Lithuania | Freshwater, shallow (~7 m), 65 km<sup>2</sup>, built in 1960, highly productive; population native | TAE, seasonal and spatial closures in 2005-2012, but only one TAC for fish all species combined; in 2012 only bycatch allowed; since 2013 commercial harvesting banned. | Minimum size limit: TL ≥45 cm; daily bag limit of 2 pikeperch (total bag ≤5 kg), seasonal closures (March 1- May 31) | Highly eutrophic, evidence for improving scientific CPUE; large recreational fishing impact (ca 110K visits in 2020) | Yes, sporadically |
| 8. Lipno Reservoir, Czech Republic | Freshwater, shallow (~7 m), 47 km<sup>2</sup>, built in 1958, highly productive; population non-native but introduced into the river system before the reservoir was built (most likely to the fishponds at the catchment in the 1950s) | All commercial harvest banned since 1976 with the exception of whitefish fishing in 1989-1996 | Minimum size limit: TL ≥45 cm; daily bag limit of 2 pikeperch (total bag ≤7 kg), seasonal closures (Jan 1- June 15) | stock collapsed during the early 2000's; currently stock appears recovering; large recreational impacts (100-150K visits a year) | Yes, ca 100,000 annually (10-15 cm long) |
| 9. Upper Lake Constance, Austria | Freshwater, 473 km<sup>2</sup>, deep (max 251 m, mean 101 m), oligotrophic. Main pikeperch habitat in the littoral area. Population introduced in late 1800s. | TAE (permits, days fishing and nets), minimum mesh size of 50 mm | Minimum size limit: TL ≥40 cm; seasonal closures (April 1 – May 30) | Over 80% of commercial and recreational catch occurs in Austria. No reliable information on population trends, aside from catch trends available. | Yes, ca 15,000 annually |

#### Commercial catches and effort

Catch data were compiled from national reports and assumed to be accurate (i.e., we did not include error estimates in commercial catches). For Galtfjärden, commercial catches for the Sea of Åland and Bothnian Sea (ICES Subdivisions (SD) 29N and 30) were used, and for Kvädöfjärden – catches from the Northern Baltic Proper (SD 27). Commercial effort and therefore commercial CPUE estimates were available for Finland (population number #1 and #2 in Table 1) and Lake Constance (#9); they were collected as number of days fishing (in #1, Heikinheimo et al. 2014), number of lifted gillnets of ≥ 40 mm mesh size (in #2, Vehanen et al. 2020) or number of commercial fisheries permits issued in Austria as reported in annual fishery commission reports (in #9, following Baer et al. 2017). Other stocks did not have sufficiently detailed estimates of commercial effort (e.g., in Lithuania only the total allowable and not the actual commercial effort is recorded) and the commercial CPUE data were not used. Commercial CPUE was standardised using general linear models as described below.

#### Recreational catches and effort

Despite being an important recreational target, for most of the stocks assessed in this study, recreational catch data were unavailable or highly uncertain. In Finland (stocks #1 and #2) recreational catches are estimates from questionnaire surveys (Heikinheimo et al. 2014). In Lithuania detailed recreational effort and catch surveys were conducted in Kaunas WR (#7) using on-site and drone-based surveys and anonymous data from a fishfinder device popular among anglers (Dainys et al. 2022b). In Czech Republic (#8) anglers are required to provide reports of all effort and catches to the Czech Angling Union, therefore there are detailed data on recreational effort and catches for the Lipno population (Vehanen at al. 2020). In Lake Constance (#9) recreational catches are the raw catches reported by anglers in log books provided with fishing licenses and compiled in annual commission reports (for 2021, see Baer and Blank 2021). For Sweden, Estonia and Curonian Lagoon in Lithuania, suitable recreational catch and effort data were not available and therefore not used.

#### Scientific surveys

Regular scientific surveys have been conducted for the past 20 –30 years in five out of the nine study sites: in Sweden, Lithuania (#3, 4, 6, 7) – using standardised gillnets; in Estonia (#5) – using trawls. Therefore, for these stocks we used standardised scientific CPUE, as described below. More details about the surveys are listed in Table S2, and for detailed information on monitoring programs see HELCOM 2019 and Vaino 2016.

#### Expert based assessment of data quality

During the process of data collection, we also asked experts from each country to evaluate the quality of recreational, commercial and scientific survey data on a 1 to 5 scale (Table S3). Full details on the ranking criteria and data quality assessment result are shown in the Supplemental Methods.

### Statistical analysis

#### CPUE estimation and standardisation

The use of CPUE as an index of relative abundance requires the removal of the effects of variation due to changes in factors other than stock biomass (such as season, location etc.). Depending on data availability, CPUE time series to be used in surplus production models (SPM) in this study have been estimated from scientific surveys, commercial or recreational catch and effort data (Table S3). To account for systematic and random variation across sampling events, we standardized CPUE using generalised linear models (GLM) with Tweedie distribution (Shono 2008), as implemented in the R packages ‘statmod’ (v. 1.4.33; Giner and Smyth 2016) and ‘tweedie’ (v. 2.3.3; Dunn 2017). Fixed effects and models varied for each stock (Table S4) and standardisation models were initiated using the full model with all explanatory variables included (Table S4), excluding those that were strongly correlated (e.g. location and depth in the Swedish coast data). The number of parameters was reduced in a backward selection manner based on Akaike’s Information Criterion (AIC; Akaike 1974). The model with lowest AIC score (cut-off at delta AIC<2, Burnham and Anderson 2002) was selected as the best model, as is commonly used in scientific CPUE standardisation routines (Forrestal et al. 2019). The parameter year was treated as an unordered factor, which means that for each year the model estimated separate coefficients. These coefficients and their estimated uncertainty values were then extracted, back-transformed to the original units and used in further analyses as standardised annual CPUE.

Due to a lack of data, we did not apply CPUE standardization to Lake Constance and Lipno reservoir stocks. In Lipno reservoir there is no commercial pikeperch harvest, and CPUE was estimated as reported annual recreational pikeperch catch divided by annual recreational effort, recorded as the number of daily angler trips per year. There was a change in angling regulations in 2009 –2015, when all natural baits (bait fishes < 20 cm TL) were banned, and subsequently only artificial lures were allowed when fishing for pikeperch. This decreased the overall angling efficiency by approximately half (expert estimate), so the total angling effort during these years was multiplied by 0.5 to reflect the relatively lower actual effort. In 2016 when natural fish bait with a minimum length of 15 cm was introduced, the efficiency of angling increased slightly, and for the years 2016 –2020 an effort multiplier of 0.75 was applied to the raw effort data. In Upper Lake Constance, CPUE was estimated as yearly commercial catches in Austria divided by the number of yearly commercial permits in the country. This approach assumes that permits reflect the amount of effort, which is unlikely to be true, but other effort or CPUE data were unavailable.

#### Surplus production models

We applied the open-source Bayesian State-Space Surplus Production Model framework, JABBA (Just Another Bayesian Biomass Assessment, Winker et al. 2018). The state-space framework enables incorporation of both observational and process errors, and the Bayesian approach allows quantifying uncertainty in the parameter estimates. We used the Schaefer type production curve (Schaefer 1954), which assumes maximum sustainable biomass at 50% of carrying capacity (also following recommendations in ICES 2022). Separate models were run for each stock using a standardised abundance index estimated above and total catches (combined commercial and recreational catches, where available). Standard error estimates associated with the CPUE index were taken from the GLM models (errors of the year parameter estimate), and catch observations were considered error free.

JABBA requires prior information on the resilience parameter *r* (intrinsic rate of population increase), carrying capacity *K* and the relative initial biomass at the beginning of the time series. Stock-specific *r* values were not available and we therefore assumed identical *r* priors for all stocks (mean = 0.25, CV=0.2) and a lognormal prior distribution. This choice was based on the classification of resilience in FishBase, where pikeperch is classified as a low resilience species with *r* values ranging between 0.05 and 0.5 (Froese et al. 2017). Priors for *K* were set as a minimum and maximum range, corresponding to 3 and 10 times of the maximum recorded catch in the time series. For the Lake Oulujärvi population, *K* priors were based on expert opinion about the history of the stock and set as 2 to 6 times the maximum catch. Priors for the initial biomass were more stock-specific and based on available knowledge about historical stock status. High relative initial biomass (0.6–0.8) was assumed for the Finnish Archipelago, Sea Galtfjärden, Kvädöfjärden, Curonian Lagoon, Kaunas Water Reservoir and Lake Constance; medium for Lake Oulujärvi and Lipno (0.2); and very low relative initial biomass was assumed for Lake Peipsi (0.05) to reflect an observed stock collapse following intensive seine netting (Saat et al. 2010). Model projections for the next 10 years were generally based on catches from the last year in the time series, as well as 50% smaller and 50% larger catches. To evaluate the model’s goodness-of-fit, we used the standard deviation of the normalized residuals (SDNR) and a runs test (Francis 2011), which assess the randomness and the magnitude of CPUE time series residuals compared to model predictions. The runs test is implemented in the JABBA package and quantitatively evaluates the randomness of CPUE fit residuals on a log-scale. A relatively good model fit is characterized by small residuals (close to zero) and a SDNR value that is close to 1.

## Results

### Data collection and quality and current stock management

#### Data quality and current assessment methods

In most countries commercial catch data reporting was judged to be of relatively high quality, typically with values of 4 or 5 (Table S3); in all cases this data was used in the final analysis. Lowest commercial catch data scores were assigned to water bodies which are shared between countries, mostly Curonian Lagoon and Lake Peipsi, both shared with Russia. Commercial effort data were considered of high quality only for the Finnish Archipelago Sea and Lake Oulujärvi. In other places, commercial effort is reported as the total allowable and not the actual effort expended (e.g., number of fishery permits in Lake Constance and allowable effort in Curonian Lagoon); is available only on much larger spatial scale (Swedish coast); or reporting requirements have changed through time (Curonian Lagoon) making comparisons problematic. As a result, commercial effort and commercial catch per unit effort (CPUE) series were only used for the Finnish Archipelago Sea, Lake Oulujärvi and Upper Lake Constance. For Upper Lake Constance commercial fishery data were not considered of high quality (only permits issued for fishing all species were recorded and not the actual effort), but it was the only source of effort data as a proxy for relative population abundance, since regular scientific surveys are not targeted at pike perch.

Fishery independent scientific surveys are conducted yearly in five out of nine stocks, they use consistent methodology and provide high quality data (scores of 4-5) (Table S3). For Lipno stock regular surveys have only started recently (since 2016). For the Finnish Archipelago Sea and Lake Oulujärvi no fishery independent scientific surveys are taking place.

The main gaps in data availability were in recreational catch and effort. Only five stocks had some estimates of recreational catches, and only one of them (Lipno) was considered of high quality and based on regular mandatory catch reports (quality of 5). For other stocks, recreational catch estimates were either limited to short time periods (Kaunas reservoir) or were based on questionnaires and therefore potentially biased (Finland) and conducted only every few years (e.g. every 5^th^ year in Lake Oulujärvi). In some stocks (Curonian Lagoon, Swedish Baltic Sea coast) there was no assessment of recreational catch whatsoever, even though recreational catches are likely to be important.

In most European pikeperch stocks studied here, current population assessment methods rely on trends in commercial catches, commercial and scientific CPUE or body size statistics (Table S1). Only two populations (Finnish Archipelago Sea and Lake Oulujärvi) have been assessed with a more complex virtual population analysis (VPA), but the assessments were conducted back in 2014 (Table S1).

#### Management policies

In most cases, management policies used for commercial pikeperch harvest are based on either total allowable catch (TAC) (two out of seven commercially harvested populations) or allowable gear numbers and types (total allowable effort or TAE) for the remaining populations. All populations have size restrictions on harvest, such as minimum mesh size and a minimum landing size, typically set at about 45 cm total length (Table 1). In Curonian Lagoon, TAC for the commercial fishery is implemented on the Russian side, while in the Lithuanian side, management since 2013 has switched to TAE restrictions only. Seasonal and spatial closures for commercial fisheries apply only in Curonian Lagoon. For two inland populations (Kaunas and Lipno reservoirs) commercial fishing is currently banned (since 2013 and 1976 respectively). Recreational fishery management typically involves daily bag limits and does not restrict total catch from the population. In addition, all stock has a minimum landing size (ranging from 40 to 46 cm), restrictions on lures (only in Lipno reservoir) and seasonal closures lasting from two to six months. Since 2009, permanent spatial closures have been implemented in Lipno reservoir and now encompass 6% of the total reservoir area; Lipno is the only pikeperch population with permanent spatial closures.

### Pikeperch stock status in Europe based on surplus production models

General linear models for CPUE standardisation showed that for most stocks, all variables included in the initial models were significant and hence retained in the final model. Based on the SDNR and runs test criteria, SPM for all but one stock produced a good fit. For these stocks MCMC chains converged, posterior distributions of estimated parameters had clear unimodal shapes and were within the original prior ranges (Fig. S1). The only stock that didn’t produce a good fit was that of Upper Lake Constance, where although the SDNR value was good at close to 1, the runs test indicated a non-random distribution of residuals. Therefore, Upper Lake Constance results should be treated with caution.

Surplus production modelling indicated that during 2019-2020 three out of nine studied stocks (Curonian Lagoon, Lipno reservoir and Galtfjärden) were in depleted state with stock spawning biomass being below the biomass at maximum sustainable yield (B_msy_) (Figs 1, 2). Four stocks (Kvädöfjärden, Lake Oulujärvi, Lake Peipsi and Upper Lake Constance) had biomass levels higher than the estimated B_msy_, although in two stocks the uncertainty ranges overlapped with B_msy_ values (Kaunas WR and Archipelago Sea). Four stocks showed strong improvements in the stock status over the last decades. Specifically, in Lake Peipsi pikeperch biomass levels were very low in the 1960-1970s, but since 1990s-2000s the stock seems to be in a good status and still improving (Fig. 2). In Lipno Reservoir, the pikeperch stock collapsed in the mid-2000s, but the population has since been recovering. The Archipelago Sea population in Finland showed a declining trend in biomass, but since 2010 the stock has stabilised, with recent signs of potential improvement. Positive trends were also seen in Lake Oulujärvi (Finland). In Lake Oulujärvi, the pikeperch population collapsed in the 1960-1970s, and despite stocking since 1985, biomass levels remained very low until the onset of natural reproduction in 1999. Since then, the population has grown rapidly and has remained relatively stable for the last decade. Positive trends in stock status were also observed in the Kvädöfjärden stock, on the eastern coast of Sweden. In contrast, three stocks (Curonian Lagoon, Galtfjärden and Kaunas water reservoir) all showed a decline in their biomass over recent decades, with Curonian Lagoon and Galtfjärden being at especially low biomass levels. In Kaunas reservoir, the declining stock appears to have stabilised, possibly as a consequence of the commercial fishing ban imposed since 2013; yet, the stock showed no clear signs of recovery. Finally, the population status in Lake Constance appeared to be good, with biomass above B_msy_ and relatively low fishing mortality. However, this result is uncertain, because the time series of CPUE and catch data didn’t include sufficient contrast in population abundance to estimate population parameters; as a result, the overall model fit is relatively poor.

**Fig. 2.**
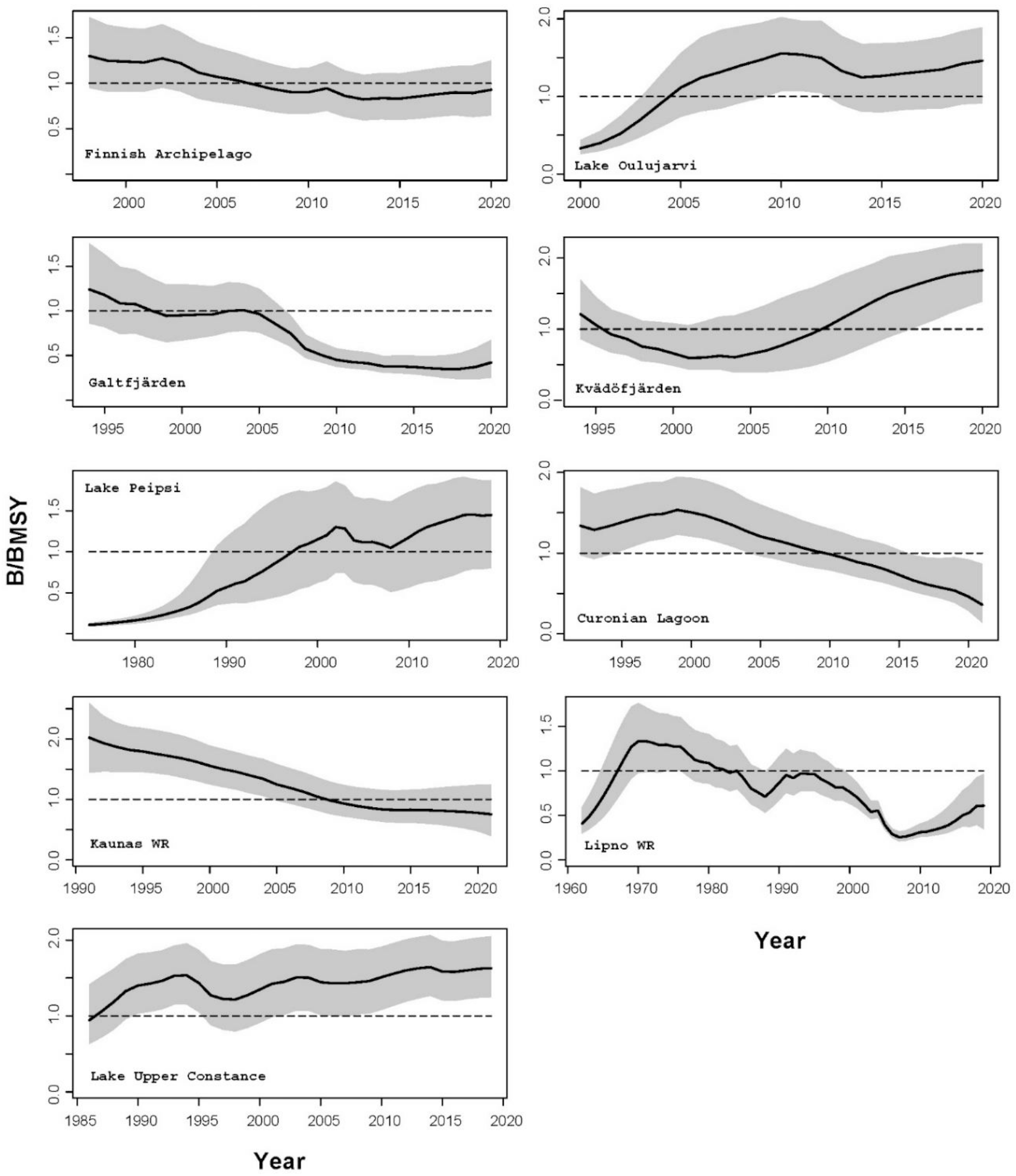
Pikeperch biomass changes in the nine studied stocks. The y axis indicates the biomass relative to the biomass at maximum sustainable yield (B_msy_). The grey area and the thick central line show the 95% and 50% posterior probabilities. Note the different time series durations for each stock, depending on data availability.

When looking at estimates of fishing mortality and projections of stock status into the future (Fig. S2), the worst scenario outcomes were for the Curonian Lagoon, Kaunas reservoir and Lipno reservoir, where estimated biomass was below the limit for maximum sustainable yield, yet commercial and recreational fishing mortality remained above the mortality required to maintain the stock at maximum sustainable yield (Fig. 1). Running model projections with different catch levels for the next 10 years indicated that if catches for these populations remained at current levels, the biomass would continue to decline (Fig. S2). Biomass projection showed that for populations to recover to B_msy_ by 2030, catches in Kaunas and Lipno reservoirs should be reduced by about half, while in the Curonian Lagoon a drastic reduction (at least 10 times) or a complete moratorium is required. Notably, for Galtfjärden stock, reported catches have been drastically decreased, and if catches remain that low, population is likely to recover in ten years. For Finnish Archipelago Sea, Lake Oulujärvi, Lake Peipsi and Kvädöfjärden current catches appear sustainable and will maintain good population status.

## Discussion

In this study we have presented an overview of data collection programs, management practices and population status and trends for nine freshwater and Baltic Sea populations of pikeperch. The studied stocks provide a good representation of northern and central European pikeperch populations across different countries, habitats, management practices and fishing impacts. Some populations are mostly exploited by commercial fisheries (e.g. in the Baltic Sea) while others are fished only recreationally (Kaunas and Lipno water reservoirs). Pikeperch is one of the most valuable commercial and recreational species and we find that many populations have some form of regular data collection programs and reasonable data quality to assess population trends. Yet, in most cases management advice is based on mean population body size, CPUE or catch trends, and regular stock assessments are rarely conducted. Our unified assessment framework indicates that three out of nine populations are in a depleted state, four populations have good status and two are at biomass levels close to B_msy_ values. Three large lake populations, in particular (Oulujärvi, Peipsi and Upper Constance) currently appear to have good stock status with biomasses mostly above B_msy_, and strong improvement over the last few decades in Oulujärvi and Peipsi. The most concerning situation was found in the two Baltic Sea stocks (Curonian Lagoon and Galtfjärden), which showed strongly declining trends over the last decade and very low current levels of biomass. Although not included in this study due to data availability, the status of Pärnu Bay (Estonia) pikeperch stock is also considered poor, due to overfishing since the end of the 1990s (Eero 2004; Eschbaum et al. 2021).

### Trends and status of Baltic Sea populations

In the coastal areas of the Baltic Sea and in the Curonian Lagoon pikeperch supports important commercial fisheries, with annual catches of 360 to 150 tons in Finland (declining trend since 2015) (LUKE 2022a), 300 t in Curonian Lagoon, and 28 t in Sweden (Fig. S3). There have been multiple concerns raised over the sustainability of these fisheries, especially in Sweden, Estonia and Lithuania. Pikeperch in the Baltic Sea has a history of being intensively exploited (Eero 2004; Mustamäki et al. 2014), as many fisheries harvest a large proportion of the pikeperch soon after or even before reaching maturity, recreational fishing pressure may be intense, and predation from great cormorants and seals have also been contributing to population decline in some areas (Mustamäki et al. 2014; Heikinheimo et al. 2015; Sundelöf et al. 2022). The status of Galtfjärden population remains poor and other studies have also shown that the mean and maximum age in the catch decreased significantly during 2002–2019 (Sundelöf et al. 2022). Interestingly, our results and earlier studies (Sundelöf et al. 2022) indicate that the Kvädöfjärden population has improved significantly over the last decade, although exact causes for this improvement are not clear. One possibility could be a decline in competition with pike, another important predator, since recent study showed that the abundance of pike has considerably decreased in Kvädöfjärden from 2005 to 2020 (Bergström et al., pers. comm.).

**Fig. 3.**
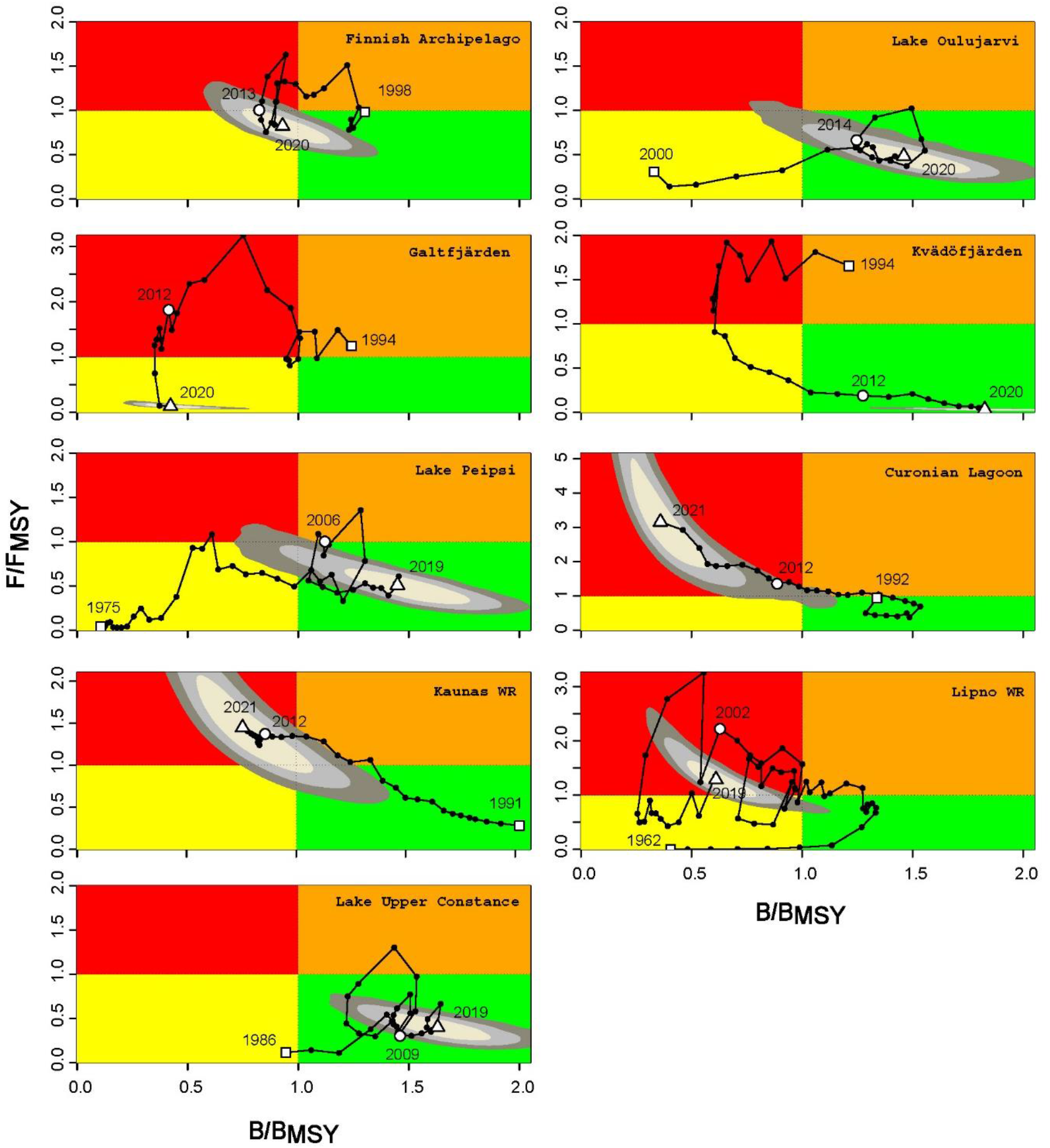
Kobe plots for the studied pikeperch stocks. The plots show the population trajectory through time (line with dots indicating stock position each year), relative to the estimated annual instantaneous fishing mortality (compared to the mortality at MSY, F/F_msy_) on the y axis and stock biomass compared to biomass at MSY (B/B_msy_) on the x axis. Red panel (top left) indicates that the stock is depleted and overfished (F > F_msy_ and B < B_msy_), the green panel (bottom right) indicates a sustainable stock status (F < F_msy_ and B > B_msy_). The orange panel (top right) indicates overfishing, but no depletion yet (F > F_msy_ and B > B_msy_), whereas the yellow panel (bottom left) shows a potentially recovering stock which is depleted but no longer overfished (F < F_msy_ and B < B_msy_). White square shows the beginning of the dataset and triangle indicates the end. The concentric grey circles show 50%, 80% and 95% posterior probability ranges.

In the Finnish Archipelago Sea the population status of pikeperch stocks appears to be relatively stable and likely improving during recent years, although population biomass is still possibly below MSY levels, given the uncertainty ranges. Improvements in the population status could be related to increased minimum landing sizes from 37 cm to 40 or 42 cm TL, depending on the area (Vainikka et al. 2017b; Olin and Raitaniemi 2021) and reductions in commercial fishing effort (LUKE 2022b). The observed increase in water temperature (Viitasalo and Bonsdorff 2022) and eutrophication are also likely improving pikeperch reproduction success and individual growth rates. Yet, recreational fishing is also considered to be an important source of mortality and recreational effort is not decreasing, although catch and release is becoming a more practice common among anglers. One of the main challenges for pikeperch management in the Archipelago Sea population is the slow progress of introducing larger mesh sizes in the commercial fishery. This is because the fishery targets a mixture of perch and pikeperch and the most used mesh sizes are at 43-45 mm, leading to a large undersized pikeperch bycatch and commensurately high mortality (Olin and Raitaniemi 2021).

Pikeperch population status in the Curonian Lagoon remains contentious, because commercial catches over the last decade appear to have remained stable (Fig. S3), yet the scientific CPUE has declined drastically over the last decade. The apparent stability of commercial catches despite declining population trends, could be explained by large increases in commercial fisheries efficiency, given that over the last decade commercial fishery adopted considerably more efficient, modified fyke nets. Yet, these fyke-nets also lead to high incidental mortality of undersized fish (Ložys et al. 2022), potentially explaining rapidly declining scientific CPUE trends. Large colonies of cormorants in the area might also have some effect on the species, however this has not been well studied for the Curonian Lagoon. Recreational catches are also likely to have large impacts yet remained unassessed. The declining population trends of the Curonian Lagoon pikeperch population have been a topic of intense discussions between Lithuania and Russia, and management bodies in Lithuania. There are plans to reduce or ban all commercial harvesting within the Lithuanian jurisdiction of the Curonian Lagoon, yet it is unclear how such a ban would affect the total population because three quarters of the Lagoon lies within Russian territory and data sharing and annual stock status negotiations ceased in 2022.

Although the importance of the commercial pikeperch fishery in the Baltic Sea has always been well recognised and actively managed, its recreational fishery remains less studied and is subject to only limited management measures. Yet, mortality for coastal fish populations from recreational fishing is likely to be high. For example, in Curonian Lagoon recreational effort is likely to be high, especially during the migration period in May-June, when part of the pikeperch population gathers in large numbers near the Klaipeda Strait connecting the Curonian lagoon with the Baltic Sea (Ložys et. al. 2017). Intense recreational fishing prior and after the seasonal closures also occurs within the delta of the Nemunas river where pikeperch migrate to spawn (Ložys et. al. 2017). Similarly, along the Baltic Sea coastline of Sweden, recreational catches can be considerably higher than commercial ones (Sundelöf et al. 2022). Recreational catch estimates are highly uncertain but in 2014 they were estimated to be in the range of 9 to 64 tonnes compared to just under 14 tonnes in the commercial fishery (Sundelöf et al. 2022). Similarly, recreational pikeperch catches in Finland generally exceed the commercial catch, although estimates again are also highly uncertain (LUKE 2022a, b). Finally, another important population in the Baltic Sea is found in Pärnu Bay, which although not included here due to limited data, used to yield substantial commercial catches (150-300 tons in the 1990s and up to 800 tons in the 1930s). During the 2000s this population collapsed, most likely due to overfishing, and has since remained in a poor state (Lehtonen et al. 1996; Eero 2004; Eesti Mereinstituut 2016). The minimum size limit has been increased from 44 to 46 cm TL and currently the main proportion of the catch is taken under ice using gillnets. It is possible that mild winters without permanent ice cover may provide an opportunity for the population to recover. However, the recreational fishery in Pärnu Bay also contributes to mortality with estimated annual catches at about 13 tonnes (Kantar 2021). Like in many places, this estimate does not include the catch of undersized fish which are released, but the incidental mortality can be high, especially during warmer seasons (Eesti Mereinstituut 2016).

Essentially, considering the poor status of some Baltic Sea populations, largely unknown recreational catches and potential impacts of seal and cormorant predation, precautionary approach would suggest increasing the minimum landing sizes for Baltic Sea populations. This is likely to eventually lead to higher yields, as the average size of pikeperch would increase (Olin and Raitaniemi 2021). Also, protection of reproduction habitats should also be among management measures (Sundblad et al. 2011). Finally, to minimise impacts of climate change, environmental fluctuations, and evolutionary trends we suggest introducing strategically located spatial closures where all commercial and recreational fishing is banned. The EU mission of Restoring our Oceans and Waters aims for 30% of land and sea to be in protected areas, which, if implemented properly, would greatly improve the resilience and status of Baltic Sea pikeperch populations.

### Origins and status of freshwater populations in lakes and reservoirs

The natural distribution of pikeperch in Europe is largely confined to coastal and brackish habitats and river deltas. Many freshwater populations in Finland, Czech Republic and Austria have been established by stocking, although many now also have natural recruitment (Table 1). For example, in Lithuania pikeperch now occurs in about 50 freshwater lakes and reservoirs (all, except for Kaunas reservoir populations were established by stocking), although only the Kaunas reservoir population had sufficient data to support quantitative assessments presented here. In Finland, pikeperch occurs in most major lakes up to the Arctic circle. It is native in about 650 lakes, but due to widespread stocking, the present distribution covers about 2300 lakes (Lappalainen and Tammi 1999). Several important native freshwater pikeperch stocks in Finland, including the Lake Oulujärvi population, collapsed during the 1960s and early 1970s (Colby and Lehtonen 1994) and have been restored through reintroductions and subsequent stocking. Due to insufficient control and planning, practically all pikeperch stocking programmes in Finland have relied on only a few source stocks, all of them of southern origin (Salminen et al. 2012). For example, the contemporary pikeperch stock in Lake Oulujärvi (64°N) stems from the Lake Vanajavesi (61°N) stock (Salminen et al. 2012) and despite effective natural reproduction in Lake Oulujärvi since 1999 (Vainikka and Hyvärinen 2012), the population is still annually enhanced by releasing about 300,000 young-of-the-year fish from multiple southern Finland populations (60-62°N). The genetic and population level impacts of such stocking could be substantial but remain largely unclear, although see Salminen et al. (2012).

In Lipno and Kaunas reservoirs some pikeperch occurred naturally before the respective damming of the Vltava and Nemunas rivers (Vostradovsky and Tichy 1999), but populations were further augmented by stocking. Regular and extensive stocking is still applied in Lipno, despite good evidence that only about 20% of adult fish derive from stocking whereas 80% have recruited from natural reproduction (microchemical analyses by Souza et al., unpublished data). A similar situation occurs in Lake Constance, where pikeperch was originally introduced in the late 1800s and fish have been stocked almost annually, despite studies showing that stocking has limited benefits for the naturally reproducing population (Löffler 1998). Stocking in Kaunas reservoir is more limited, but still occurs almost annually. Lake Peipsi in Estonia represents unusual exception in that the population is natural and is not supplemented by stocking.

Most of the lake populations, except for Lipno Reservoir, have been harvested commercially, although commercial fishing has also been banned in Kaunas Reservoir since 2013. Recreational fishing surveys suggest that in many large Finnish lakes, including Lake Oulujärvi, and in Lake Peipsi in Estonia commercial fishing is considerably more important than recreational catches. For other populations, especially those occurring in more densely populated areas, recreational catches represent the major source of mortality. Nevertheless, recreational fishing effort and catch data for most lake and coastal populations remains too spatially or temporally unresolved to assess potential impacts because catches are often estimated only through country-wide surveys or licence sales. As a result, despite the importance of pikeperch for both commercial and recreational fishing, targeted management often remains limited to size restrictions and sometimes daily bag limits.

### Pikeperch population status and environmental factors

Commercial and recreational harvests seem to be among the main factors affecting pikeperch abundance in both coastal and freshwater populations (Lehtonen et al. 1996; Mustamäki et al. 2014; Olsson 2019). Nevertheless, environmental factors undoubtedly also play a role. Firstly, nutrient runoff and consequent eutrophication, affect both adjacent freshwater bodies and coastal areas of the Baltic Sea. In general, pikeperch are believed to benefit from eutrophication and rising water temperatures, because turbid, warm water positively influences pikeperch spawning success and increases somatic growth rates (Kjellman et al. 2003; Lappalainen et al. 2003; Lehtonen et al. 1996; Pekcan-Hekim et al. 2011; Veneranta et al. 2011). For example, improvements in land and water usage and increased water clarity in the Czech Republic during recent decades appear to have had negative impacts on pikeperch stocks (Tesfaye et al., submitted). Recent analyses of lake pikeperch populations in Lithuania show large increases in mean population body size over the last decade, likely driven by both the ban on commercial fishing (since 2013), as well as increasing temperatures and eutrophication (Audzijonyte et al., unpubl.).

Climate change and warming are also likely having positive impacts on pikeperch, as this species benefits from warm water conditions. This is especially true in Lake Oulujärvi and populations across Northern Finland, where pikeperch occurs close to its northern range limit and recruitment is strongly correlated with summer temperatures. It is likely that warming conditions are also leading to faster juvenile growth rates with potentially earlier maturation at smaller body sizes, as is expected for many fish under the ‘temperature size rule’ (e.g., Audzijonyte et al. 2016). However, for most populations, data on maturation and growth rates are not collected routinely, hence trends in reproduction with size and their ecological implications remain unassessed. Generally, faster growth rates should potentially improve population status, and if maturation does occur at smaller body sizes fixed minimum size limits may mean that populations are recruiting to fisheries at older ages. Yet, life-history changes will make it harder to assess long-term population trends, as many data poor assessment methods generally assume that the underlying population parameters remain unchanged.

One potentially important factor limiting pikeperch populations is a lack food availability for juveniles, especially during overwintering months. As proportions of predatory fish in populations decline due to intensive harvesting (Myers et al. 2003; Fogliarini et al. 2021), cyprinids, such as bream and roach, often become the dominant species in freshwater and brackish coastal systems. For example, the biomass of roach in Kaunas Reservoir has nearly quadrupled after the commercial fishing ban, as this species is not regularly targeted by anglers (Dainys et al. 2022a). Both young pikeperch and cyprinids compete for food in their early life, yet when the abundance of animal food is low, especially during winter, cyprinids can switch to feeding on detritus and periphyton (Richeux et al. 1992), whereas pikeperch cannot. Intraspecific competition between juveniles in strong years classes has also been suggested to contribute to make pikeperch recruitment density dependence (Lappalainen et al. 2009). Finally, another factor that is likely to affect some pikeperch populations is growing predation pressure from seals and cormorants in the Baltic Sea. Mustamäki et al. (2014) found that cormorant predation may have contributed to a decrease observed in small pikeperch abundance, as during some years cormorant consumption was almost equal to the commercial catch. In the Archipelago Sea the average mortality of 2-4 year old pikeperch by cormorants ranged at 5 to 30% of the total estimated annual mortality (Heikinheimo et al. 2015).

### Data-limited methods for stock assessment

The majority of freshwater and coastal fish populations are considered data limited, in that regular ageing data are not collected and this prevents applications of age-structured stock assessment models (Costello et al. 2012). However, such stocks can still have reasonable amounts of alternative data, including commercial or recreational catches and effort, and length frequencies, collected during regular monitoring or by recreational anglers. This provides an opportunity to develop and apply a more unified assessment framework and to obtain a better picture of stock status. Surplus production models provide a suitable alternative and, due to improved statistical methodology and user-friendly software interfaces are increasing in popularity (Cousido – Rocha et al. 2022). Despite their simplicity, SPMs are widely used for assessing data-limited stocks (Wang et al. 2014; Cousido-Rocha et al. 2022) and are among the simplest of full stock assessment tools which can inform managers about reference points, stock size and recommended harvest rates. In some cases, they can be more robust than parameter rich age-structured models (Ludwig and Walters 1985, 1989). It is important to note that SPMs make a key assumption that population abundance is driven by catches, population regeneration or growth rate (*r*), and carrying capacity (*K*). This means that all aspects of production are pooled into one production function *r* ignoring age, size, and sex structure. Similarly, environmental variation is either ignored or incorporated into process uncertainty, such as in case of JABBA, although environmental effects can be deterministic and if sufficient data are available should ideally be partitioned from random error (Deyle et al. 2018). Moreover, SPMs perform best when time series of catch and population abundance data have sufficient contrast to estimate population growth parameters. If data lack contrast, then this may suggest that catches have always been managed at sustainable levels or that the stock may be more influenced by factors other than catch e.g., environmental variation (Haddon 2020). This is likely to be the case for Upper Lake Constance, where pikeperch is not an important target species and population abundance is likely to be regulated by other processes.

### Conclusions and future research needs

Our study suggests that overall pikeperch population status and trends in Europe are reasonably healthy, with only two out of nine populations in this study showing strong depletion without signs of recovery. This is encouraging, considering the ecological and economic importance of pikeperch in Europe. That pikeperch also benefits from warm and eutrophic conditions, implies the current environmental trends of warming and nutrient load are likely to further improve population status. Yet, fishing mortality imposed on many populations remains high and is likely to be even higher than estimates provided in this study, if recreational catches were fully accounted for. As such, monitoring and management of recreational fishing appears to be one of the main priorities for pikeperch stock sustainability.

Despite growing recognition that recreational fishing may have large impacts on freshwater and coastal populations, recreational fishing reporting and management remains limited throughout the world. Most countries do not require anglers to report their catches, with a single notable exception being the Czech Republic, where all angling trips must be reported. Yet even here, it is not possible to identify species specific effort, because anglers do not have to report their target species, and they do not have to report released catches. For all stocks, recreational catches, if regulated, are controlled with a daily bag limit and minimum landing sizes, yet total catch is not controlled and levels of compliance are mostly unknown. Although proportions of fish being released during angling trips is increasing (Arlinghaus et al. 2002; Cooke and Cowx 2004; Bartholomew and Bohnsack 2005), depending on species, size, and handling, post-release mortality can be as high as 90 % (Muoneke and Childress 1994; Saas and Shaw 2020), which means that catch and release angling is still likely contributing to population mortality. Maximum size limits are only implemented for coastal fish populations in Sweden, despite strong evidence that larger and older females of many fish species have better spawning success than younger ones (Berkeley et al. 2004; Olin et al. 2012). Permanent spatial closures that ban all fishing (marine or freshwater strictly protected areas) are still very rare, despite their demonstrated importance for population resilience.

Finally, with rapid climate and other environmental changes and increasing popularity of recreational fishing, it is important that pikeperch populations are regularly monitored with consistent and comparable scientific methodology and that the scientific community further improves its data sharing and collaboration on population status assessments. This was the goal of our study, and we hope that this effort will continue into the future.

## Data availability

Raw data from scientific, commercial and recreational catches and effort can be requested from respective institutions and authors. Standardised CPUE effort and catch times series used in this analysis, together with the R codes are available on https://github.com/astaaudzi/EuropeanPikeperch

## Acknowledgements

This study has received funding from European Regional Development Fund (Project No. 01.2.2-LMT-K-718-02-0006) under grant agreement with the Research Council of Lithuania (LMTLT). In Sweden data curation and analyses were supported by the Swedish Agency for Marine and Water Management (Förvaltningsmål nationellt förvaltade arter för år 2020-2022” (HaV dnr 896-20) and the Czech National Agency of Agricultural Research, project QK22020134 Innovative fisheries management of a large reservoir. Data collection for Lake Oulujärvi has been funded by Fortum Power and Heat Ltd., UMP-Kymmene Ltd., city of Kajaani and the municipality of Paltamo, and data management by H2020-INFRAIA (#730938, 871120). The authors thank Freddie Heather and Romain Forestier for help with figures.

## Supplementary Information

**Table S1.** Previously applied stock assessment methods for the pikeperch stocks studied.

| Country, institution | Water body (stock) | Time period for current stock analysis approach and periodicity of assessment | Stock assessment methods used |
| --- | --- | --- | --- |
| Finland, Luke | **1. Finnish Archipelago** | 1980-2019: annual | VPA (Heikinheimo et al. 2014)<br>Based on commercial and recreational catch data and age samples (on average 2000 per year), weight samples to covert weight catches to numeric catches (Heikinheimo et al. 2014) |
| Finland | **2. Lake Oulujärvi** | 1992-2020: annual | VPA (Vainikka and Hyvärinen 2012)<br>CPUE (temporal trends in CPUE), Mean size, fish specific measurements (length, age, growth).<br>Proportion of mature pikeperch |
| Sweden, SLU Aqua | **3. Galtfjärden**<br>Swedish coastal area | 2003-2015 | CPUE GES Catch-curve, CPUE (temporal trends in CPUE), Proportion of mature pikeperch (Mustamäki and Mattila 2015) |
| Sweden, SLU<br>Aqua | **4.Kvädöfjärden**<br>Swedish<br>coast/Archipelago<br>Sea (3 sites) | 1995-2008 | Commercial CPUE, coastal fish monitoring<br>CPUE, mortality (Mustamäki et al. 2014) |
| Estonia,<br>institution | **5. Lake Peipsi** | 2000-2019: annual | Biomass and abundance estimation from scientific<br>trawl survey, swept area method |
| Lithuania,<br>Nature Research<br>Centre | **6. Curonian Lagoon** | 1992-2021: annual | Mean size, CPUE (temporal trends in CPUE), fish<br>specific measurements (length, weight), surplus<br>production models (since 2019) |
| Lithuania,<br>Nature Research<br>Centre | **7. Kaunas WR** | 1997-2021: annual | Mean size, CPUE (temporal trends in CPUE), fish<br>specific measurements (length, weight), surplus<br>production models (since 2020) |
| Czech Rp,<br>BCAS | **8. Lipno reservoir** | 1990-2020 | CPUE (temporal trends in CPUE), fish specific<br>measurements (length, weight, age) acoustic<br>surveys and fry community surveys available for<br>some years. |
| Germany,<br>Switzerland,<br>Austria | **9. Lake Constance** | None | No formal stock assessment or informal<br>management with CPUE or other indices is<br>performed. Commercial catch is recorded by<br>country: annually since 1910, monthly since 1975<br>(catch is mostly incidental - not focal fish);<br>angler catch recorded annually by country since<br>1978 (targeted by few anglers); very limited fish<br>specific measurements (length, age) |

**Table S2.**
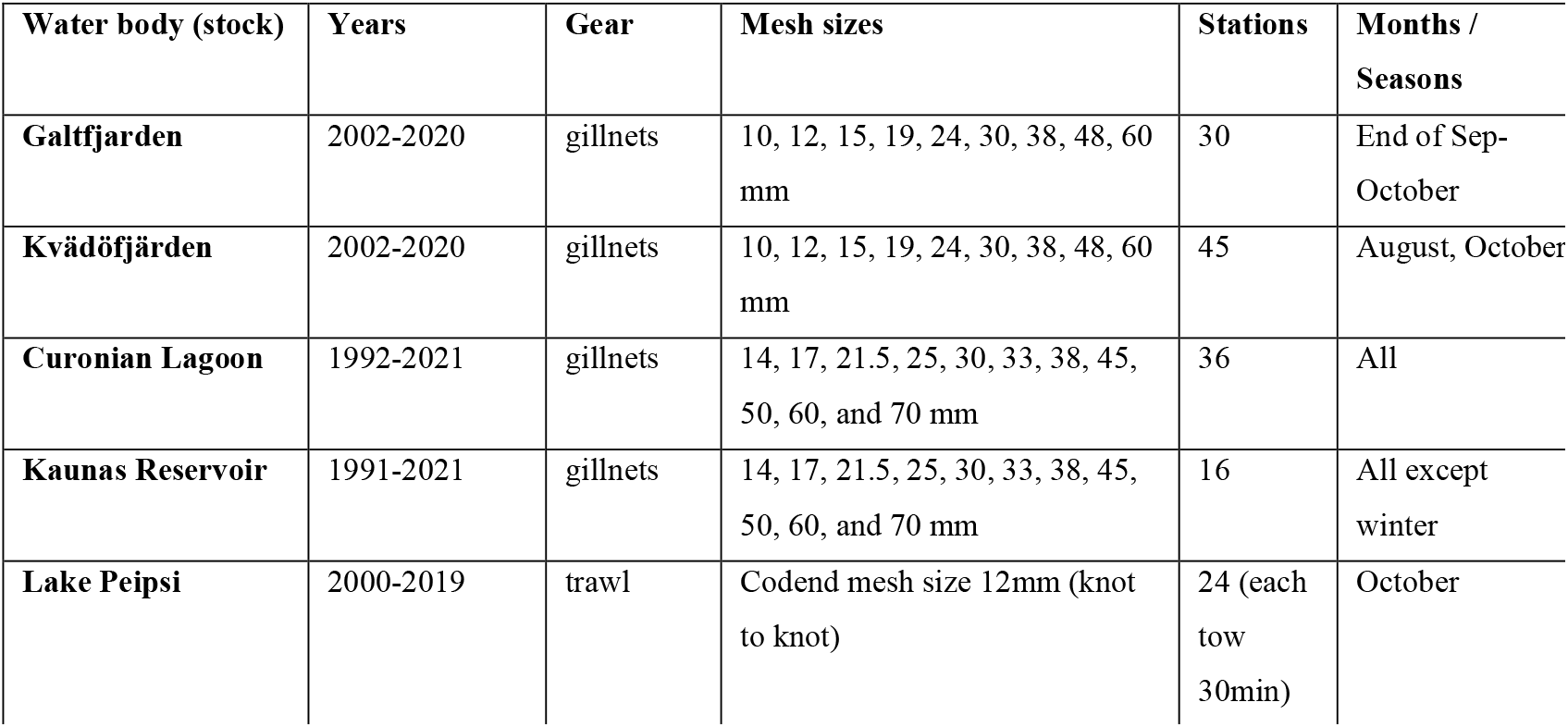
Details of scientific survey collection methods for data used in this study.

### Supplemental Methods

For commercial catches, the score of 5 meant highly reliable commercial catch or effort data from the assessed area, 4 – reliable data, but from a larger or smaller area and possibly not reflecting assessed area catches, 3 – some misreporting expected, 2 – extensive misreporting expected, 1 – extensive misreporting and large temporal or spatial data gaps. For commercial effort, the score of 5 indicates consistent and reliable reporting of actual effort by gear and fishing date, 4 – consistent reporting of actual effort, but where gears or dates are pooled (into seasons or months) and small levels of misreporting could be expected, 3 – effort data reported but requirements have changed through time and some misreporting is expected, 2 – effort data reported, extensive misreporting, large data gaps, 1 – only very general effort data (e.g. number of vessels or gears operating in the area), effort from different spatial area, changes in requirements, large data gaps. For recreational catch data the score of 5 – regular assessment of catches or effort based on creel surveys or other direct evidence, 4 – regular assessment on catches or effort based on questionnaires or other surveys, 3 – occasional assessment of catches and/or effort and extrapolation to other years, 2 – catch or effort estimation based on licence sales or similar information, 1 – expert opinion and anecdotal evidence. Likewise, for recreational effort data the score of 5 indicates regular assessment of effort based on creel surveys or other direct evidence (mandatory reporting of all trips, including zero catch trips), 4 – regular assessment of effort based on questionnaires, other surveys or mandatory reporting of non-zero catch trips, 3 – occasional assessment of effort with large temporal gaps, requiring extrapolations, 2 – effort estimation based on licence sales or similar general information, 1 – expert opinion and anecdotal evidence. For scientific surveys the value of 5 indicates yearly surveys and consistent methodology and proper spatial and temporal coverage, 4 – yearly surveys and consistent methodology but limited spatial coverage, 3 – sporadic surveys with temporal gaps in data collection, 2 and 1 – sporadic surveys and inconsistent methodology.

**Table S3.** Scientific survey, commercial and recreational catch and effort data available for the studied stocks and data used in this study (**in bold**). For each data source the time series and data quality assessment (in parenthesis) are shown. Com – commercial, Rec – recreational. NA – indicates that no data was available.

| Water body and stock | Fishery independent scientific surveys | Catches |  | Effort |  |
| --- | --- | --- | --- | --- | --- |
|  |  | Com | Rec | Com | Rec |
| **1. Finnish Archipelago** | NA | **1998-2020 (5)** | **1998-2020 (4)** | **1998-2020 (4)** | NA |
| **2. Lake Oulujärvi** | NA | (since 1974)<br>**2000-2020 (4)** | (since 1995)<br>**2000-2020 (4)** | (since 1974)<br>**2000-2020 (5)** | NA |
| **3. Galtfjärden** | **2002-2020 (5)** | **1994-2020 (4)** | NA | 2009, 2011-2021 (3) | NA |
| **4. Kvädöfjärden** | **2002-2020 (5)** | **1994-2020 (4)** | NA | 2009, 2011-2021 (3) | NA |
| **5. Lake Peipsi** | **2000-2019 (5)** | (since 1931)<br>**1975 – 2019 (3)** | (3) | (5) | NA |
| **6. Curonian Lagoon** | **1992-2021 (5)** | (since 1947)<br>**1992-2021 (3)** | NA | 1992-2021 (3) | NA |
| **7. Kaunas Reservoir** | **1991-2021 (5)** | **1991-2012 (4)** | **1991-2021 (3)** | NA | 2020 (5) |
| **8. Lipno** | 2003, 2008-12, 2016-21 (3) | Banned | **1991-2019 (5)** | NA | **1991-2019 (5)** |
| **9. Upper Lake Constance** | NA | **1986-2019 (3)** | **1986-2019 (2)** | **1986-2019 (1)** | NA |

**Table S4.** Fixed effects in statistical general linear models used for CPUE standardization. The table shows the full list of factors in the initial model. The final models were determined based on Akaike information criterion and factors that were deemed insignificant and removed from final models are shown *in italics*.

| Water body (stock) | Abundance index (CPUE) | Model used for CPUE standardization |
| --- | --- | --- |
| Finnish Archipelago | commercial | Year + month + location + effort + gear |
| Lake Oulujärvi | commercial | Year + month + location + no. gillnets |
| Galtfjärden | scientific | Year + location |
| Kvädöfjärden | scientific | Year + location |
| Curonian Lagoon | scientific | Year + season + location + gillnet material + mesh size + soak time + gillnet length |
| Kaunas WR | scientific | Year + season + location + gillnet material + *mesh size* + soak time + gillnet length |
| Lake Peipsi | scientific | Year + age |

### Figures

**Fig. S1** Prior and posterior distributions of the population parameters (r, K and psi) estimated in JABBA.

See priors and posteriors for each population at the end of this document

**Fig. S2:**
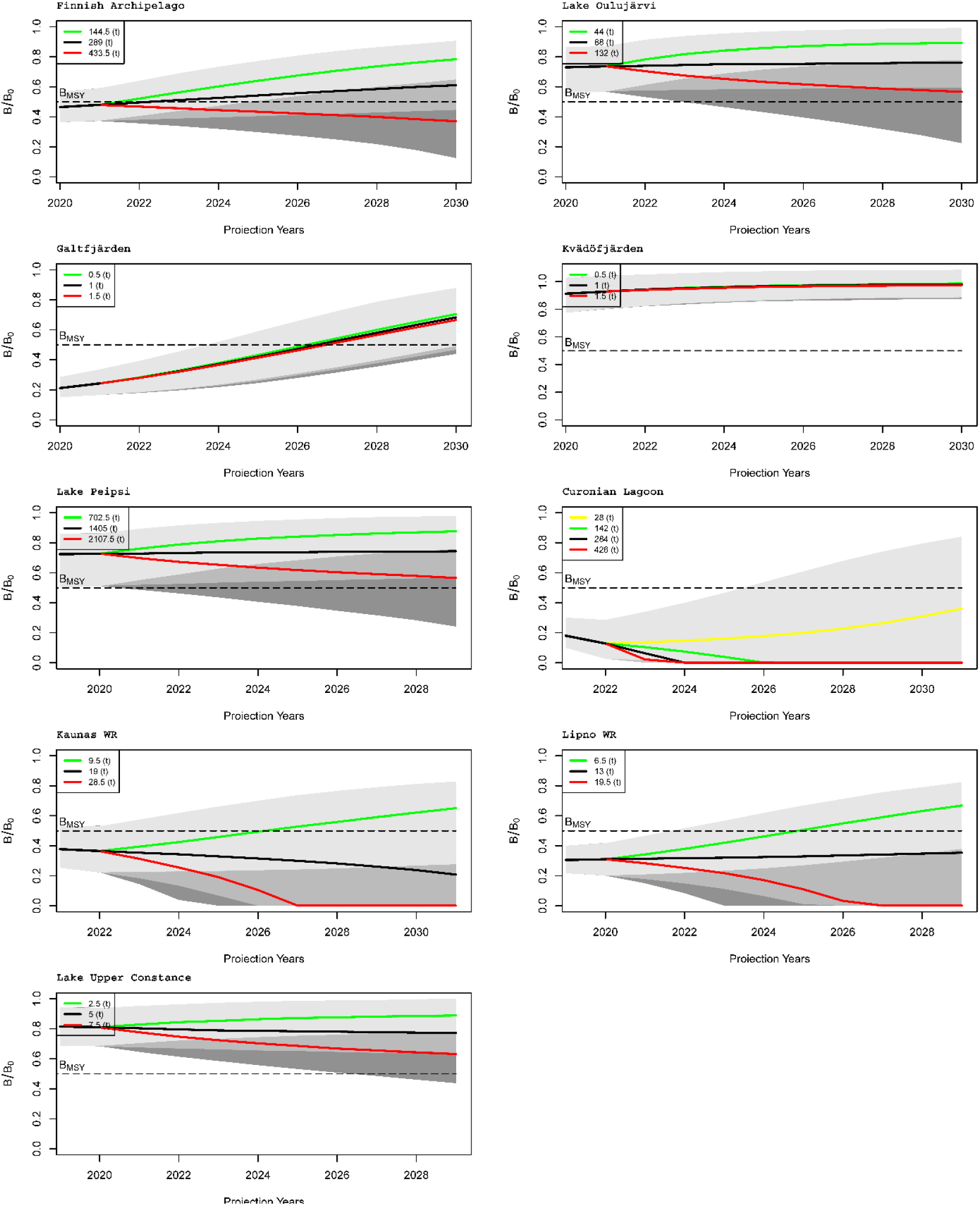
Projections of pikeperch biomass if a current catch prevails (black line), with 50% lower catches (green line), and with 50% higher catches (red line). Yellow line shows 10 times lower catch (in the Curonian Lagoon).

**Fig. S3:**
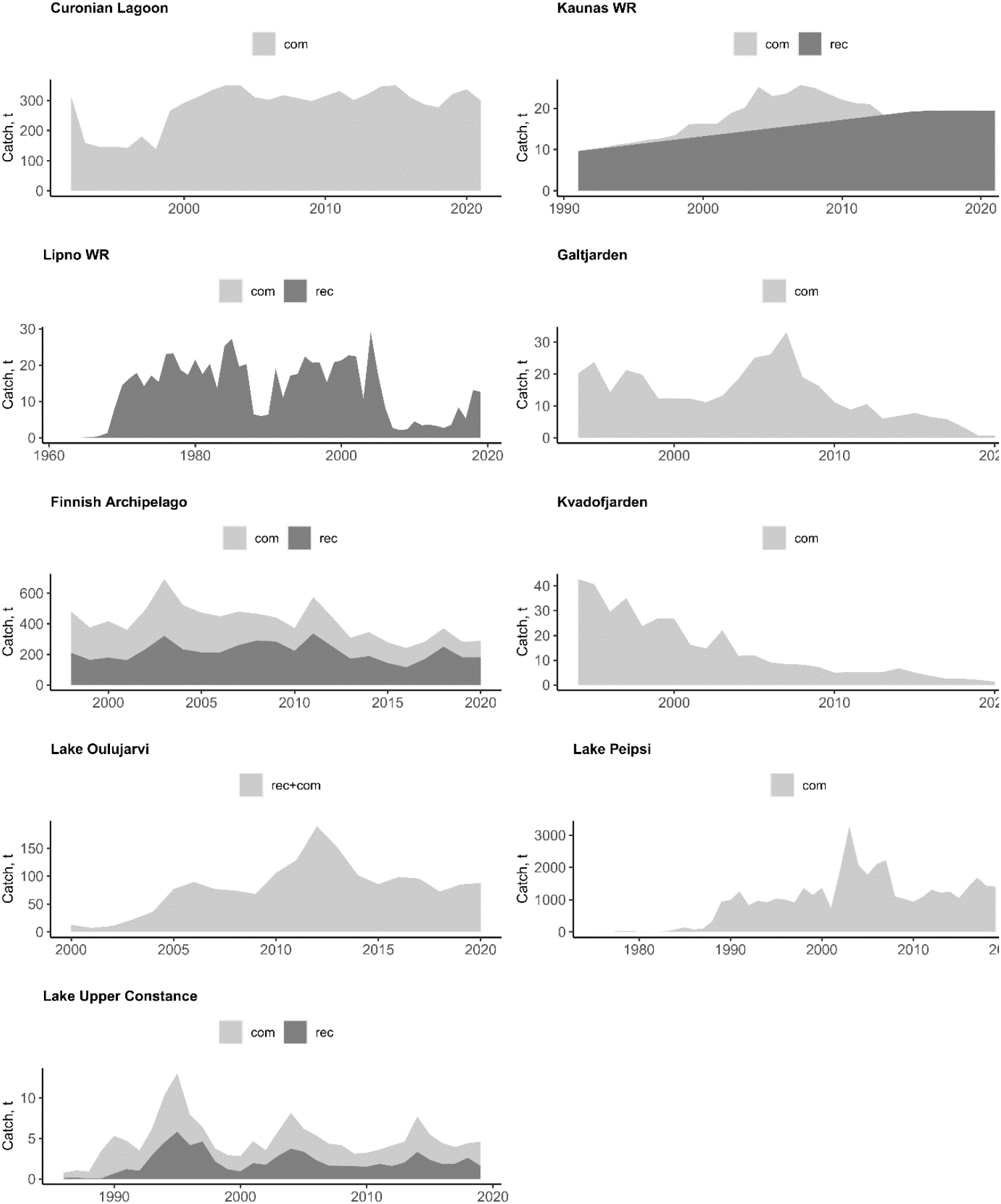
Pikeperch **c**atches in the areas studied (com - commercial, rec - recreational). In Lake Oulujärvi, only total catches are available (rec+com). In Kaunas WR the assumption has been made that recreational catches doubled since 90s (see Dainys et al. 2022).

---

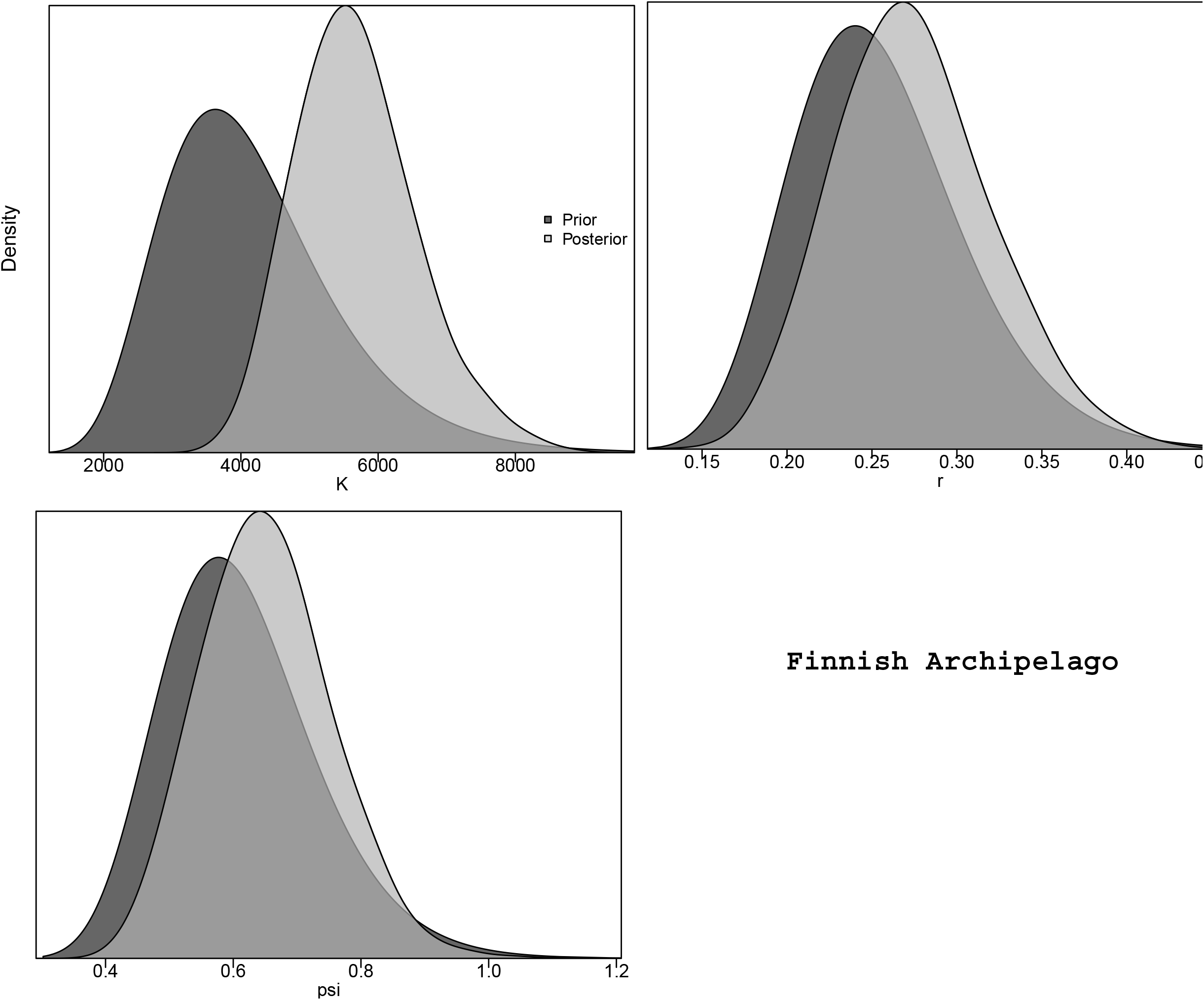

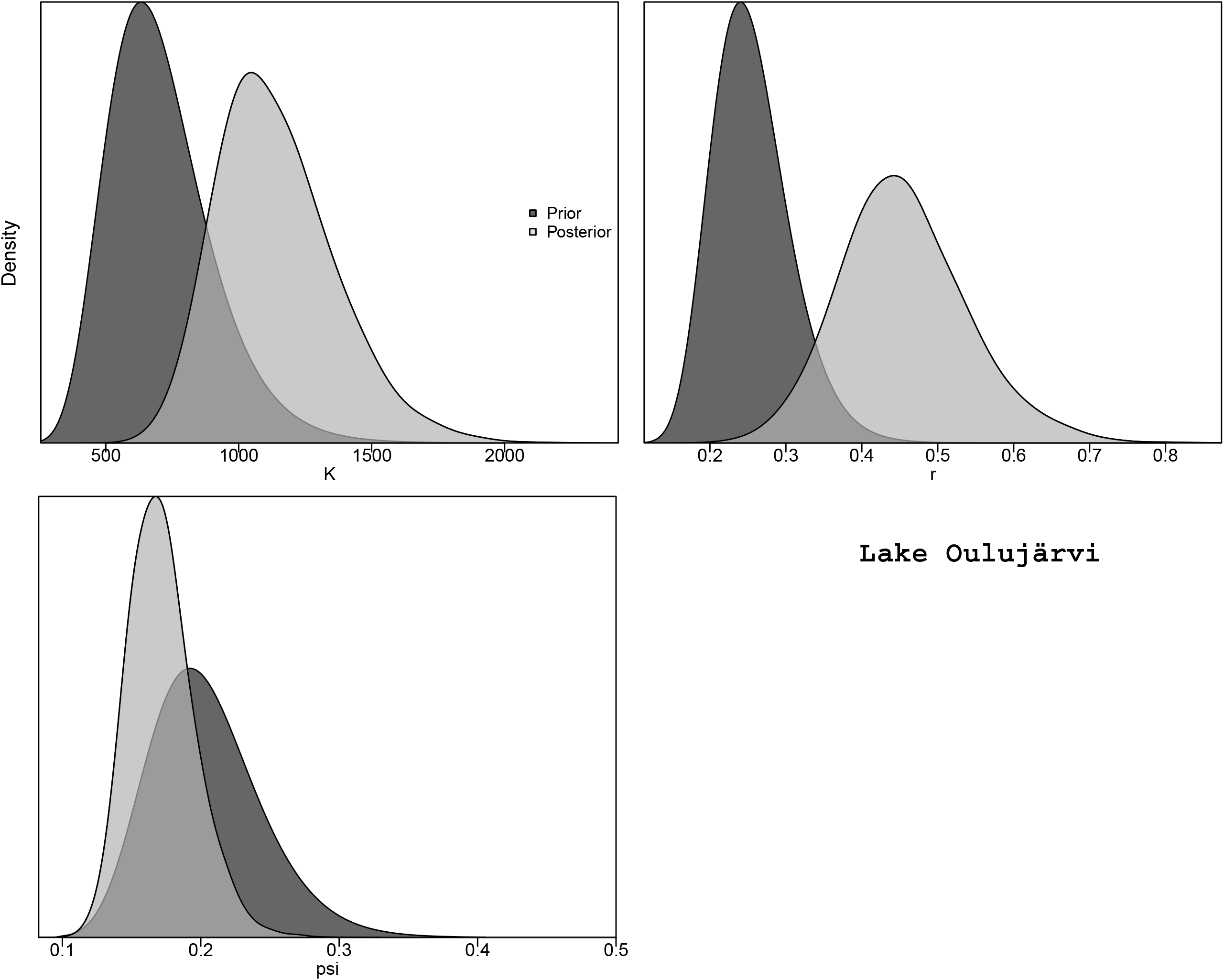

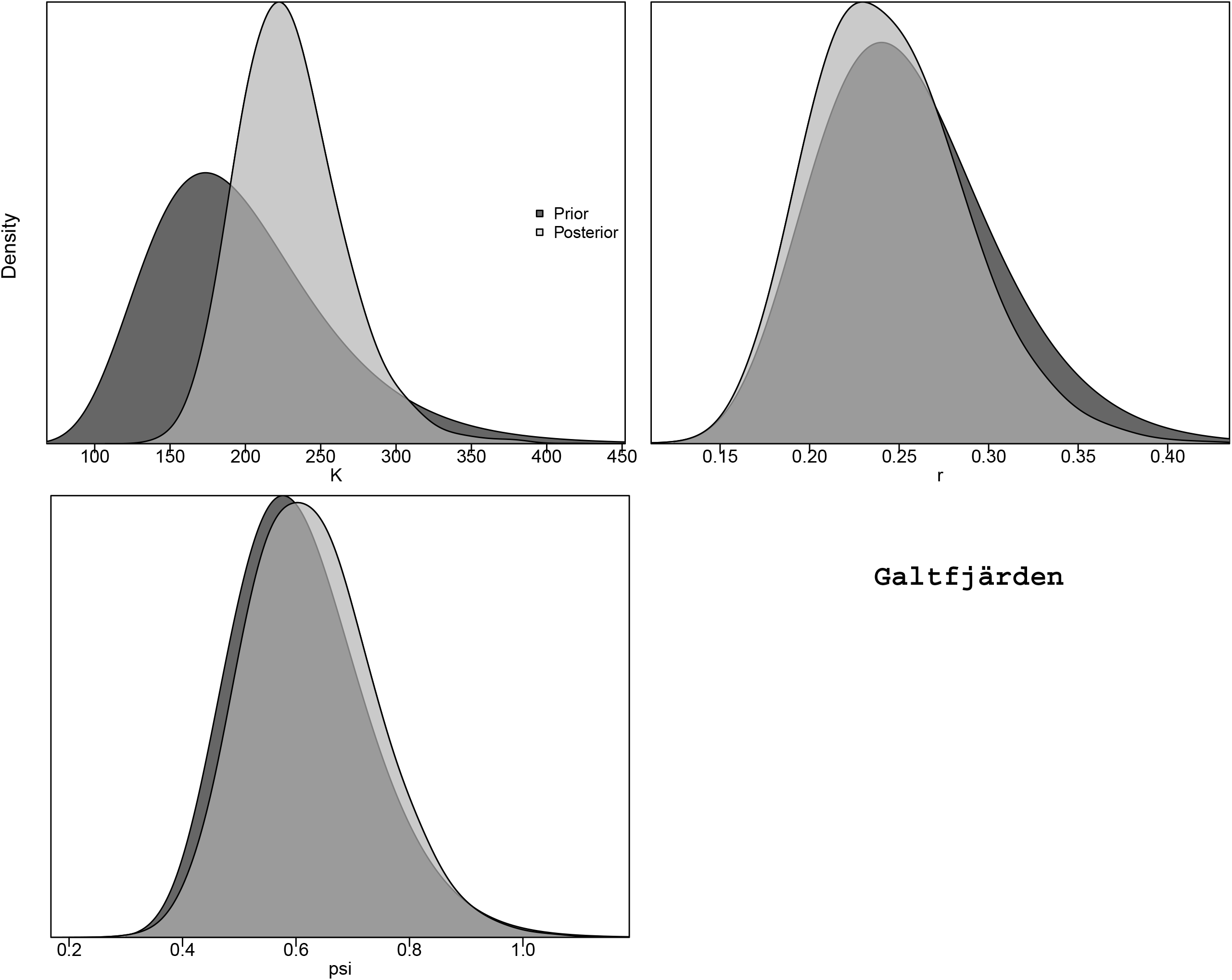

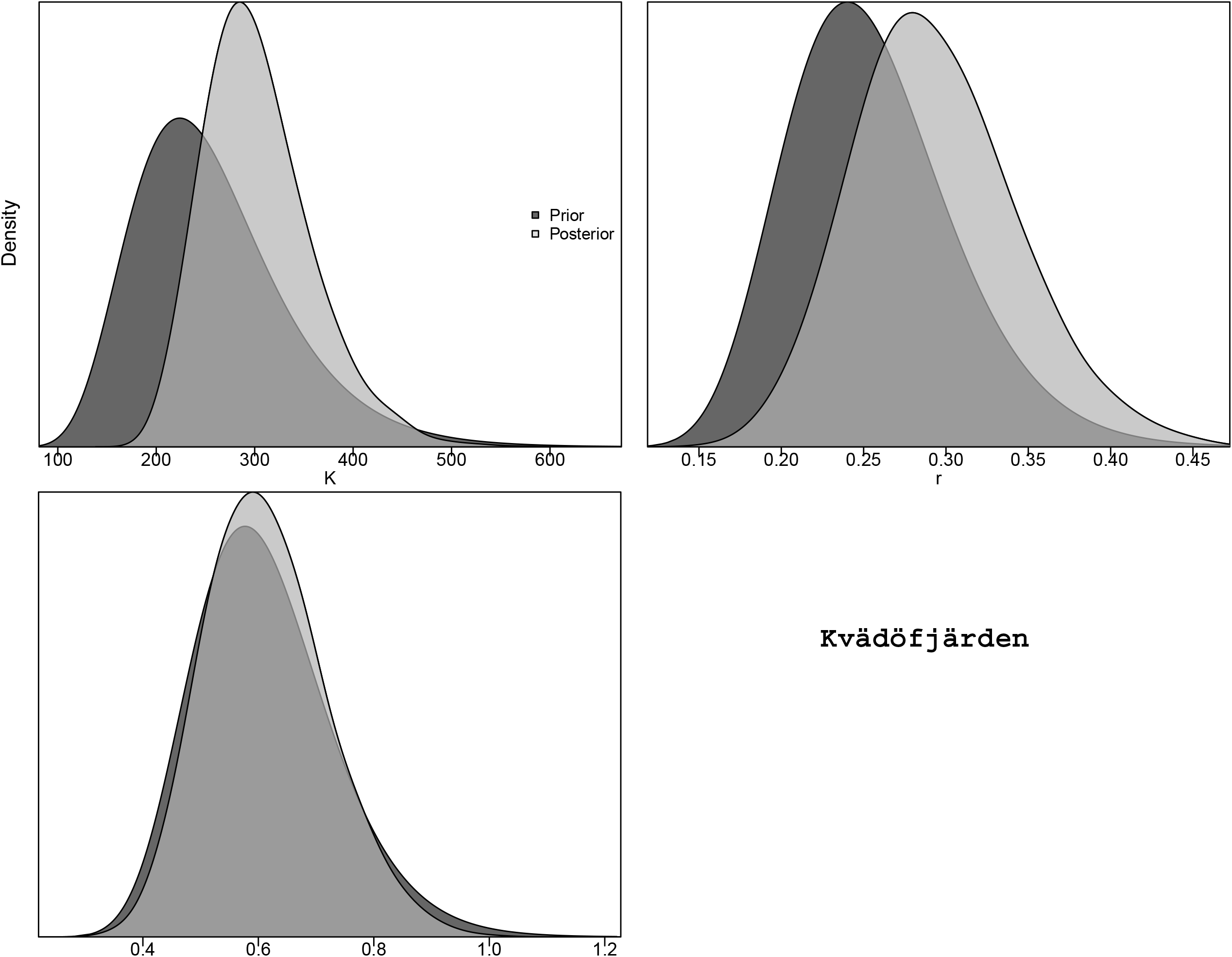

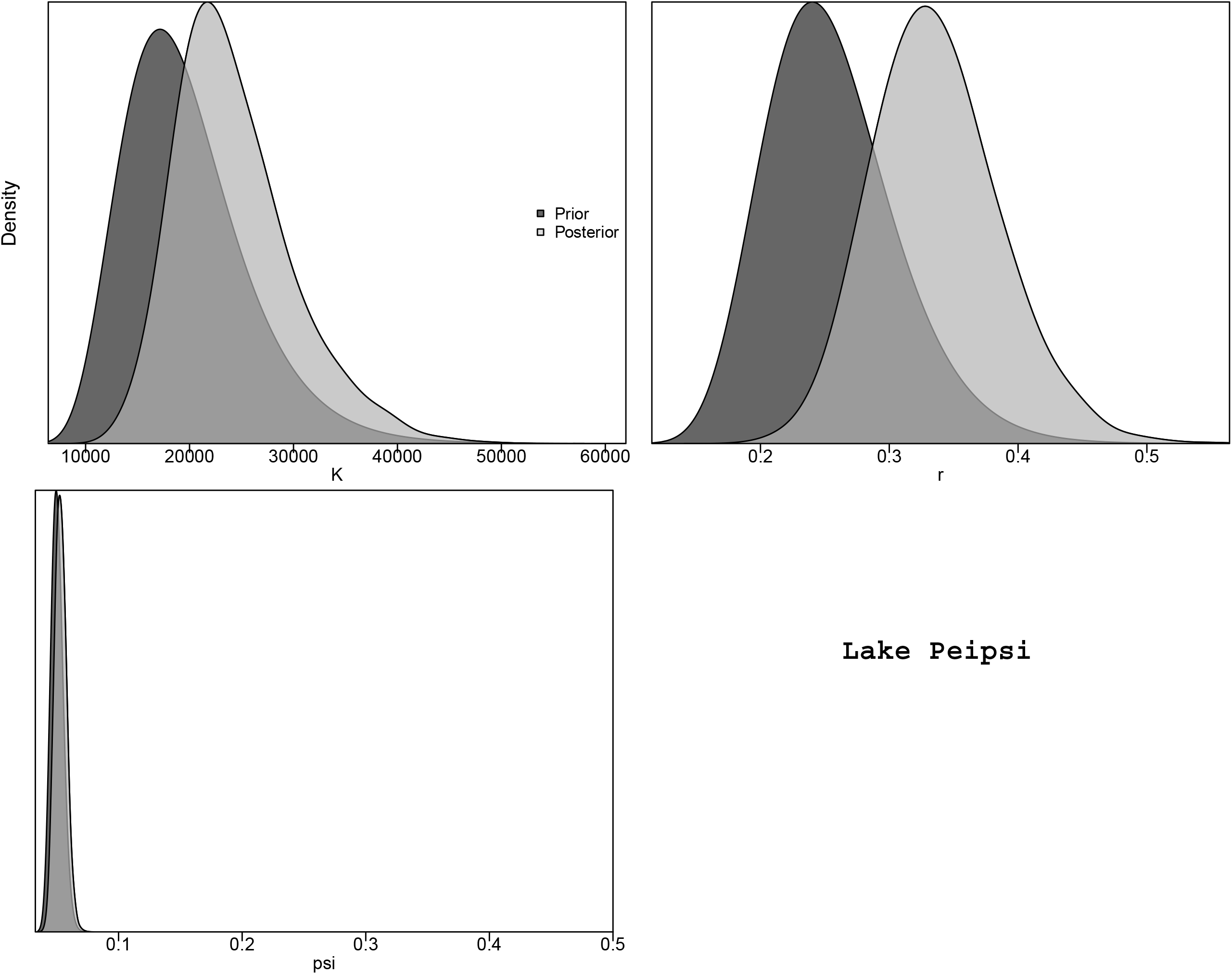

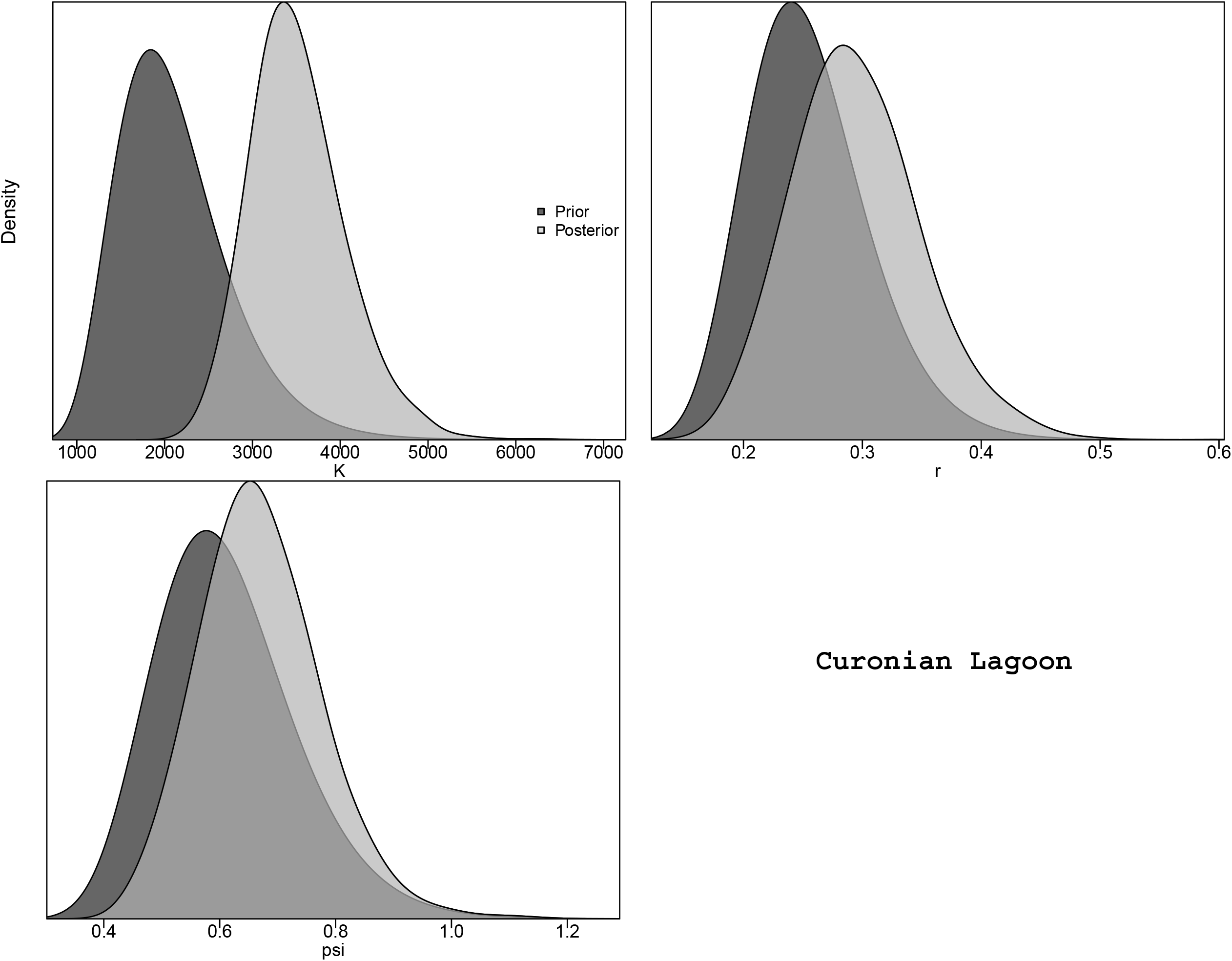

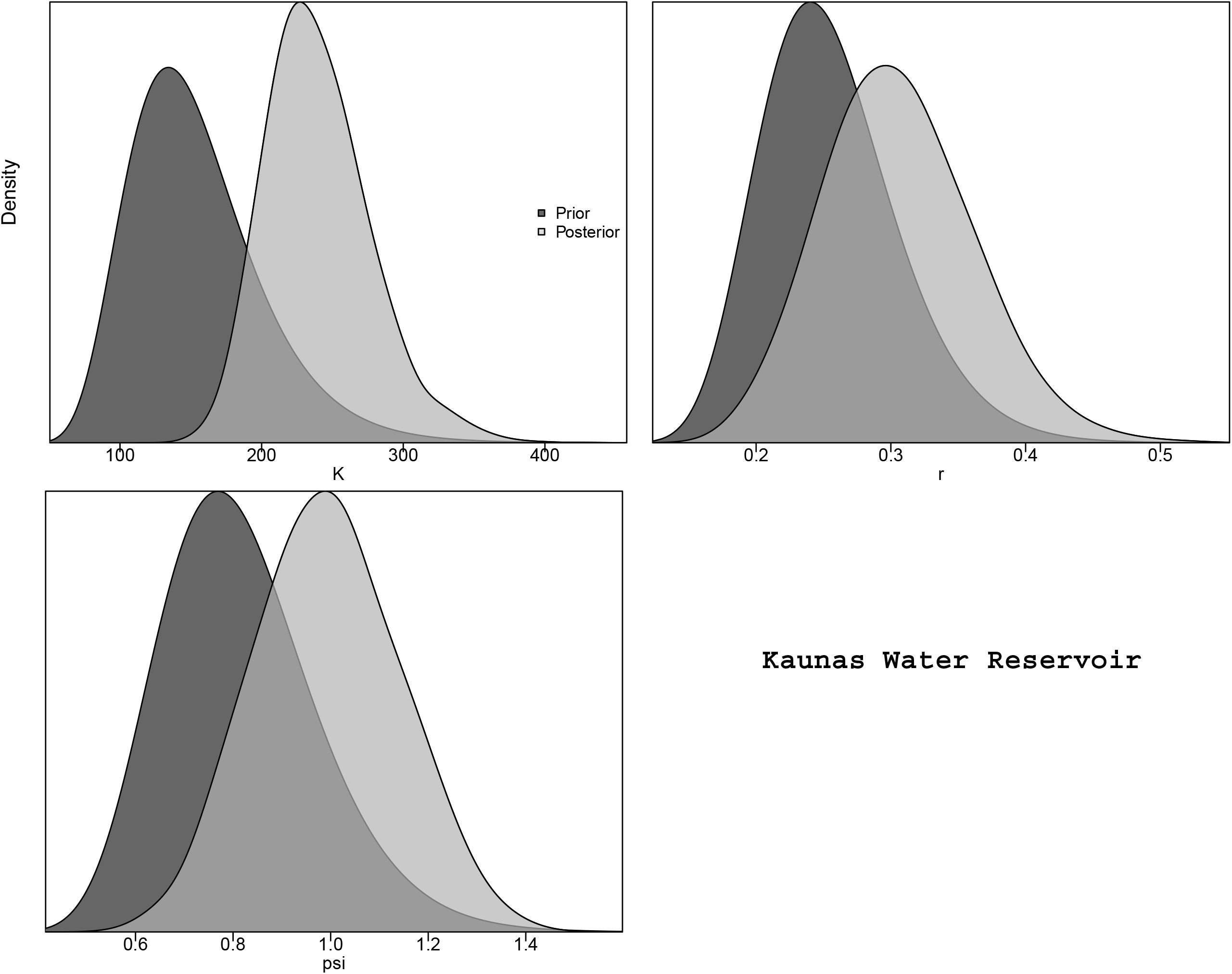

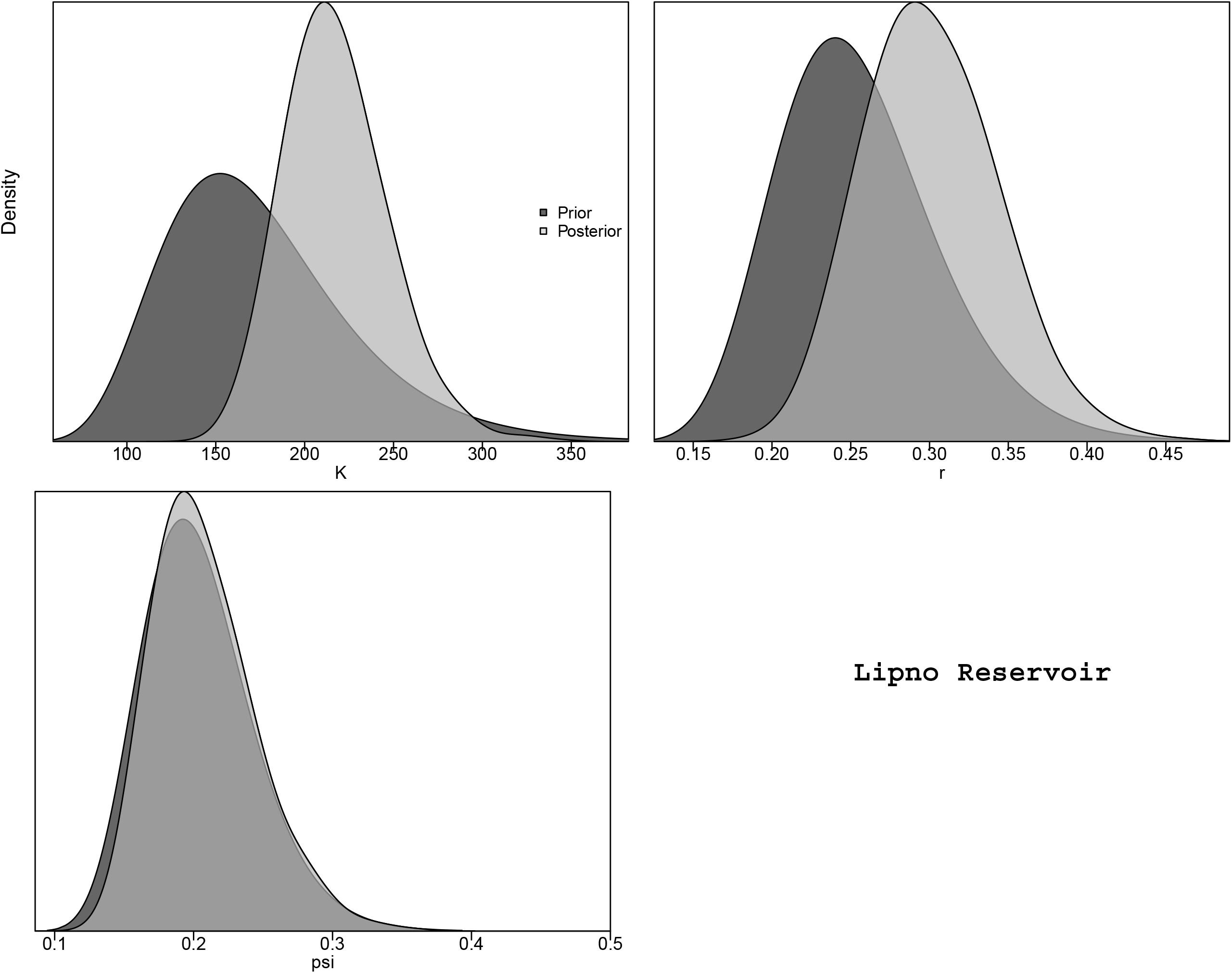

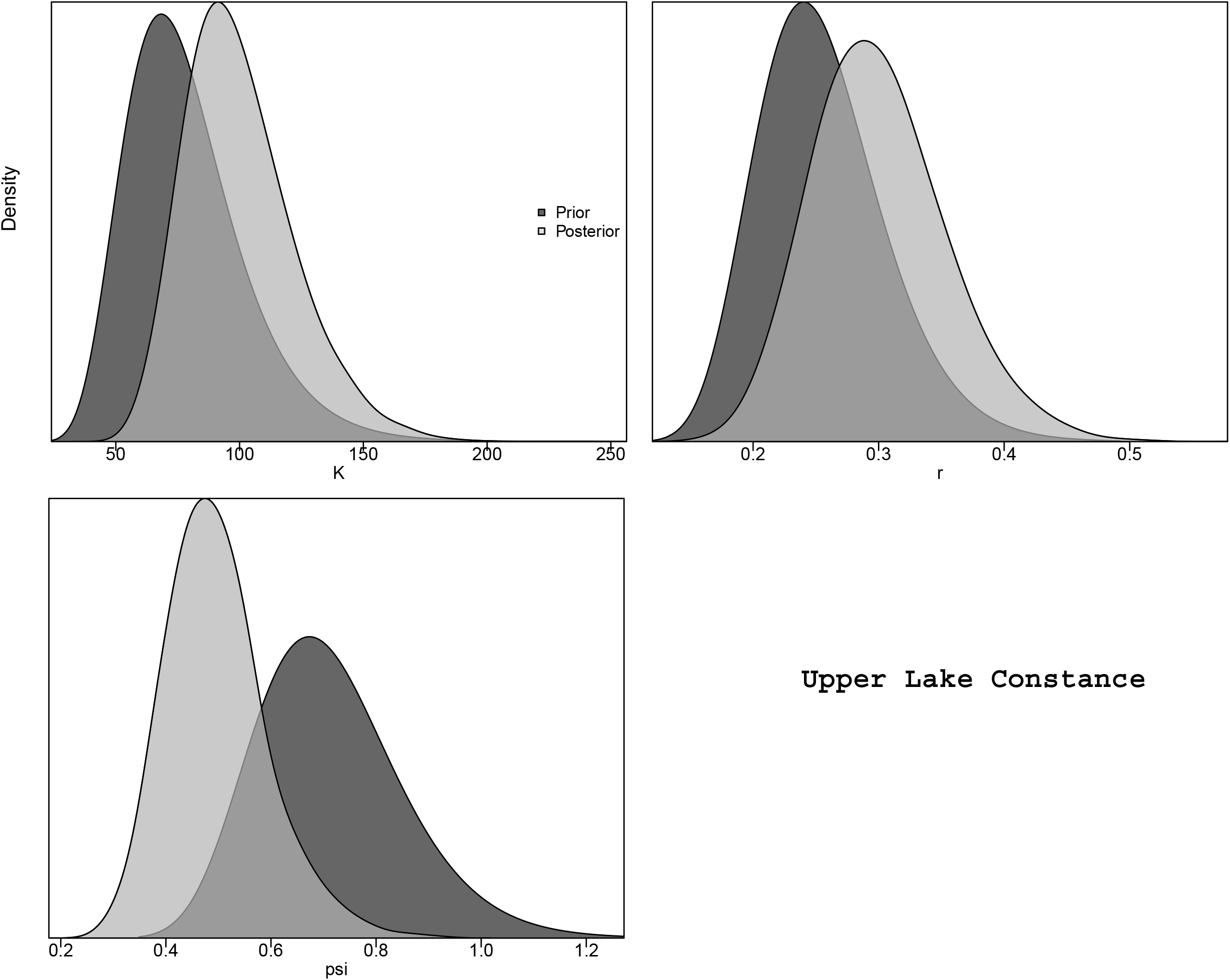

## Notes

https://github.com/astaaudzi/EuropeanPikeperch

### Competing Interest Statement

The authors have declared no competing interest.

